# ETFL: A formulation for flux balance models accounting for expression, thermodynamics, and resource allocation constraints

**DOI:** 10.1101/590992

**Authors:** Pierre Salvy, Vassily Hatzimanikatis

## Abstract

Since the introduction of metabolic models and flux balance analysis (FBA) in systems biology, several attempts have been made to add expression data. However, directly accounting for enzyme and mRNA production in the mathematical programming formulation is challenging because of macromolecules, which introduces a bilinear term in the mass-balance equations that become harder to solve than linear formulations like FBA. Furthermore, there have been no attempts to include thermodynamic constraints in these formulations, which would yield an even more complex mixed-integer non-linear problem.

We propose here a new framework, called Expression and Thermodynamics Flux (ETFL), as a new ME-model implementation. ETFL is a top-down model formulation, from metabolism to RNA synthesis, that simulates thermodynamic-compliant intracellular fluxes as well as enzyme and mRNA concentration levels. The formulation results in a mixed-integer linear problem (MILP) that enables both relative and absolute metabolite, protein, and mRNA concentration integration. The proposed formulation is compatible with mainstream MILP solvers and does not require a non-linear solver. It also accounts for growth-dependent parameters, such as relative protein or mRNA content.

We present here the formulation of ETFL along with its validation using results obtained from a well-characterized *E. coli* model. We show that ETFL is able to reproduce proteome-limited growth, which FBA cannot. We also subject it to different analyses, including the prediction of feasible mRNA and enzyme concentrations in the cell, and propose ETFL-based adaptations of other common FBA-based procedures.

The software is available on our public repository at https://github.com/EPFL-LCSB/etfl.

**Author summary:** Metabolic modeling is a useful tool for biochemists who want to tweak biological networks for the direct expression of key products, such as biofuels, specialty chemicals, or drug candidates. To provide more accurate models, several attempts have been made to account for protein expression and growth-dependent parameters, key components of biological networks, though this is computationally challenging, especially when also attempting to include thermodynamics. To the best of our knowledge, there is no published methods integrating these three types of constraints in one model. We propose here a transparent mathematical formulation to model both expression and metabolism of a cell, along with a reformulation that allows a computationally tractable inclusion of growth-dependent parameters and thermodynamics. We demonstrate good performance using community-standard software, and propose ways to adapt classical modeling studies to expression-enabled models. The incorporation of thermodynamics and growth-dependent variables provide a finer modeling of expression because they eliminate thermodynamically unfeasible solutions and consider phenotypic differences in different growth regimens, which are key for accurate modeling.

## Introduction

Metabolic modeling, which helps makes sense of the metabolism in a biological network, is an important tool for engineering biocatalysts, with applications in biofuels, drug design, microbial community analysis, and personalized medicine. Model accuracy is instrumental to the success of these applications through an efficient engineering of the host organisms. However, incorporating expression information into metabolic networks poses a significant challenge, and most current models do not even attempt it—effectively excluding an important network in biological systems that can drastically affect results. In metabolic engineering, strains are modified and controlled at the genome level through the transcriptome, and the effects are observed at the fluxome level, which accounts for the range of metabolic reactions in an organism. In between these two levels is the proteome that actually performs the biochemical transformations according to the genetic template, though it is this middle step in the process that cannot yet be robustly and efficiently incorporated into models of metabolic systems. Because of the complex interplay between these different layers of control, understanding expression and incorporating this into future models is key for improving metabolic engineering.

Classically, model-based strain design has relied on a tool that uses the DNA sequence of an organism to detail the network of metabolic reactions that happen inside a cell of that organism, which is called a genome-scale model (GEM). With current technologies and tools like metagenome sequencing [1], it is possible to generate GEMs for hundreds of different species at a time. GEMs are particularly amenable to flux balance analysis (FBA), which models metabolism at the fluxome level using linear optimization techniques. However, plain FBA has been known to predict biochemically unrealistic solutions like free high-flux cycles or thermodynamically infeasible pathways. It also scales growth linearly with carbon uptake, which is not observed at high-uptake fluxes. FBA also fails to capture growth-dependent and protein-level effects, such as enzyme saturation or proteome-related limitations. Hence, several efforts have been made to supplement FBA with additional constraints to improve its predictive power. For example, thermodynamics-based flux analysis (TFA) [2, 3] uses thermodynamic constraints to enforce thermodynamically consistent reaction directionalities and to allow the integration of metabolomics. Resource balance models add a total proteome capacity constraint, as formulated in Beg *et al.* [4], to model the proteome-related limitations of the cell, as enzymes have to compete for the constrained total amount of cellular proteins. Frameworks like GECKO [5] further build on this resource balance idea and include flux constraints based on proteomics, such as *v ≤ V*_*max*_ = *k*_*cat*_ [*E*] as well as a constraint on the total proteome mass. Finally, metabolomics and expression models (ME-models) [6, 7] were the first to integrate the entirety of the expression mechanisms of the cell from the bottom-up, including mRNA and protein synthesis.

However, simultaneously accounting for all of these constraints is challenging because of the formulation of each method, as TFA models involve integer variables that yield a mixed-integer linear program (MILP), whereas ME-models involve bilinear constraints that require special optimization procedures and a solver [8–10]. Mixing these methods would require the inclusion of integers in ME-models, which is not straightforward and would lead to more complex mixed-integer non-linear programs (MINLP) that are computationally intensive to solve. Additionally, the amount of RNA and protein, the RNA and protein expression rates, and their stabilities are all growth dependent [11], and including accurate representations of these variables leads to even more complex, non-linear models. Meanwhile, although resource balance models such as GECKO could theoretically be integrated into TFA or ME-models in the current formulations, to the best of our knowledge, no link with TFA or ME-models has been proposed. Therefore, the metabolic engineering community needs a common formulation for these methodologies to build the most accurate models.

We investigated the development of such a framework and propose herein a unified formulation for Expression and Thermodynamics-enabled FLux models (ETFL) that can account for the above integration issues. In ETFL, we address the compatibility of the formulations by expressing the growth rate variable in bilinear products as a piece-wise constant function. This reformulation allows us to transform the problem into a MILP, which we can solve efficiently using widely available open source or commercial solvers. The resulting model is then effectively able to directly integrate thermodynamic constraints as well as expression constraints and growth-dependent parameters. In this model, metabolite, enzyme, and mRNA concentration levels are explicitly defined to enable fast and easy omics integration. Finally, we show an application of this framework to a well characterized *E. coli* model, iJO1366 [12].

## 1 Results and Discussion

### 1.1 Formulation of the expression problem

To transparently account for expression mechanisms and increase the predictive power of our models, we needed to derive the equations that could bridge the biochemistry with the optimization problem that is ETFL. Here we present a summary of these equations, and detail their derivation in the section Materials and Methods. We derived these equations using assumptions similar to those used in the formulation of the GECKO [5] and ME-model [6–8].

This formulation relies on derivations rooted in the biological mechanism of expression and depends on a number of biochemical parameters related to the cell. In particular, the mass balances of the macromolecules are expressed using concentration variables. Each mass balance will yield an equation where the concentrations of the macromolecules will be variables, thus effectively formulating a new constraint of the model and allowing us to calculate concentration values by solving the model.

We can write the quasi-steady state mass balance for macromolecules as follows:

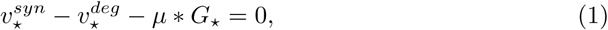

where ⋆ represents the indexing of the macromolecule, 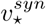 is the synthesis term, 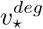 is the degradation term, and *µ ∗ G*_⋆_ is the dilution term. The detail of the derivation is available in the Materials and Methods.

Using this formalism, for each macromolecule we can define and link together a synthesis flux, a degradation flux, and the macromolecule’s concentration. Knowing enzyme concentrations allows us to bound the variables representing metabolic reaction fluxes with their maximum catalytic rate according to the classical equation:

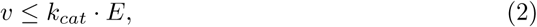

where *k*_*cat*_ is the catalytic rate constant of the enzyme *E* with respect to flux *v*. In this same fashion, we can also constrain the synthesis flux for the peptides, which are then assembled into enzymes. Peptide synthesis is simply a metabolic reaction that consumes energy (under the form of GTP) and charged tRNAs and produces a peptide and uncharged tRNAs. The catalytic rate of the reaction is proportional to the maximum ribosomal catalytic rate divided by the length of the peptide to be synthesized. The same can be said about mRNA synthesis, which uses nucleoside triphosphates and is catalyzed by the RNA polymerase. The details of these constraints are explained in the Materials and Methods.

The part of the matrix that has been added to the FBA problem to account for expression has been termed the expression problem (EP). Although this initial formulation is bilinear, we will detail in the following section how we cast it to a MILP.

#### 1.1.1 Biomass reaction synthesis and mass balance

In FBA, the biomass reaction is an artificial, lumped reaction that represents the consumption of metabolites in proportion to the cell growth rate. This consumption reflects nucleoside triphosphate (NTP) requirements for mRNAs, amino acid requirements for proteins, lipid requirements for the cell wall, or metal ion needs. Biomass reaction inclusiveness depends on the modeling assumptions made during the model curation process and can vary significantly among models of the same species. The consumed amount of each metabolite is usually estimated experimentally by measuring the the amounts of these metabolites in dried cell mass. Because the stoichiometric ratios of metabolites in the biomass reaction are fixed, the abundance of metabolites is the same for all growth rates. This simplifying assumption, necessary in FBA, goes against experimental evidence. Neidhardt and Curtis [11] report for instance that mRNA and protein mass ratios in the cell change with growth rate.

Because ETFL has explicit expression requirements through transcription, translation, and tRNA-charging reactions, it is possible to account for varying ratios of NTPs and amino acids as the growth rate changes, an effect that is captured in experiments [11]. In this context, the approximation made in FBA can be written using ETFL terms:

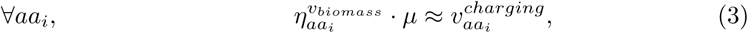

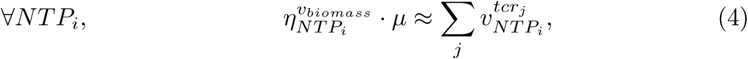

where *v*^*biomass*^ represents the biomass equation, and 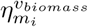 is the expression-related participation of metabolite *m*_*i*_ in the biomass reaction. Hence, to avoid accounting for the expression requirements twice (once through the biomass equation, once through the EP), it is necessary to remove the participation of these metabolites linked to expression from the biomass reaction.

#### 1.1.2 Summary

Here we show the formulation of the constraints of ETFL. For clarity, we use different indexing sets, each referring to a specific object in the model, detailed in Table 1. The variables are detailed in Table 2, and the parameters are in Table 3

**Table 1.** Indices used in the formulation.

| Index letter | Indexed variables |
| --- | --- |
| $i$ | Metabolite |
| $j$ | Reaction/Flux/Enzyme |
| $l$ | Gene/Peptide/mRNA |
| $s$ | Binary coefficient for growth discretization |
| $u$ | Binary coefficient for interpolation discretization |

**Table 2.** Variables used in the formulation.

| Symbol | Variable |
| --- | --- |
| $\mu$ | Growth rate |
| $v_j^\pm$ | $j^{th}$ net positive/negative biochemical flux |
| $E_j$ | Concentration of the $j^{th}$ enzyme |
| $F_l$ | Concentration of the $l^{th}$ mRNA |
| $P_l$ | Concentration of the RNA polymerase assigned to the $l^{th}$ mRNA |
| $R_l$ | Concentration of the ribosome assigned to the $l^{th}$ peptide |
| $T_{aa_i}^u$ | Concentration of the $i^{th}$ uncharged tRNA |
| $T_{aa_i}^c$ | Concentration of the $i^{th}$ charged tRNA |
| $v_l^{tsl}$ | Translation rate of the $l^{th}$ gene |
| $v_l^{tcr}$ | Transcription rate of the $l^{th}$ gene |
| $v_j^{asm}$ | Assembly rate of the $j^{th}$ enzyme |
| $v_j^{deg}$ | Degradation rate of the $j^{th}$ enzyme |
| $v_l^{deg}$ | Degradation rate of the $l^{th}$ mRNA |
| $v_{aa_i}^{charging}$ | Charging rate of the $i^{th}$ tRNA |

**Table 3.** Parameters used in the formulation.

| Symbol | Parameter |
| --- | --- |
| $k_{cat}^{j,\pm}$ | Forward/backward catalytic rate constant of the $j^{th}$ net biochemical flux |
| $k_{deg}^j$ | Degradation rate constant of the $j^{th}$ enzyme |
| $k_{deg}^l$ | Degradation rate constant of the $l^{th}$ mRNA |
| $\eta_l^j$ | Stoichiometry of the $l^{th}$ peptide in the $j^{th}$ enzyme |
| $\eta_{aa_i}^l$ | Stoichiometry of the $i^{th}$ amino acid in the $l^{th}$ peptide |
| $L_l^{aa}$ | Length in amino acids (aa) of the $l^{th}$ peptide |
| $L_l^{nt}$ | Length in nucleotides (nt) of the $l^{th}$ mRNA |
| $L_{rib}^{nt}$ | Ribosome footprint size on mRNA, in nucleotides |
| $\rho$ | Ribosome occupancy |
| $\pi$ | RNA polymerase occupancy |

*Catalytic constraints*

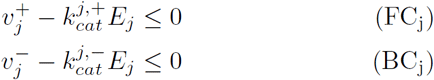

*Metabolite mass balance*

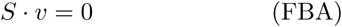

*Expression mass balance*

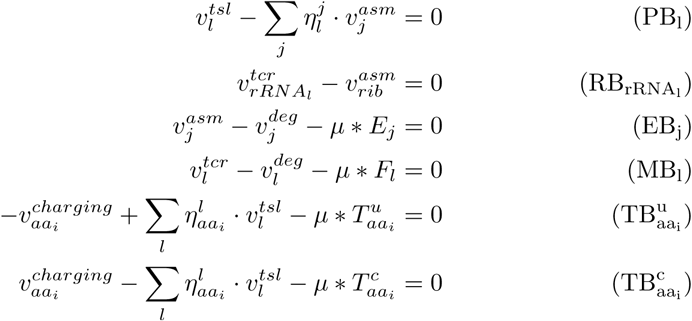

*Degradation fluxes*

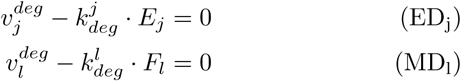

*Expression constraints*

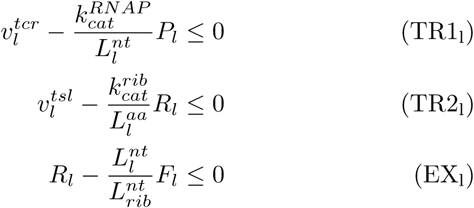

*Total capacity*

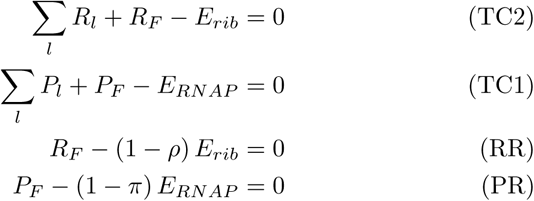

### 1.2 Reformulation

#### 1.2.1 Bilinearity of the problem

The main issue with the EP formulation presented above lies in the continuous bilinear terms that describe the dilution of the macromolecules,

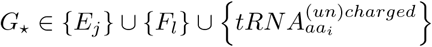. We use ⋆ as a placeholder for the indexing of *G*. Using previous notations for the synthesis, degradation, and growth rate:

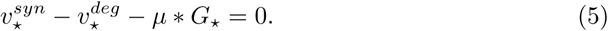

In this state, the dilution term is bilinear, and the formulation requires a bilinear solver or potentially a mixed-integer bilinear solver if thermodynamics are to be added. The original ME-model formulation has similar terms as we are presenting here [6, 7]. As such, its recent adaptation in Lloyd *et al.* [8] uses the two-level algorithm SolveME [9] that requires a dedicated non-linear solver. We present instead a MILP approximation of the problem that makes it compatible and solvable with mainstream MILP solvers. We achieve this through the discretization and linearization of the bilinear products. This operation can be understood as locally approximating the bilinear problem by several linear subproblems and choosing the best approximation.

#### 1.2.2 Approximation of the growth rate

In ETFL, we approximate the growth rate *µ* in bilinear products with a piecewise-constant function 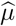 (0^th^ order approximation) to transform the continuous bilinear terms into mixed (integer × continuous) bilinear terms. This simplifies the problem, as these mixed bilinear terms can be linearized in a MILP setting using the Petersen linearization scheme [13], a particular case of the Glover linearization scheme [14] that was previously used in metabolic engineering by Hatzimanikatis *et al.* [15, 16].

Let 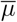 be an upper bound to *µ*, (*p, N*) *∈ N* ^2^, *p ≤ N*. We can approximate *µ* with the following 0^th^ order approximation:

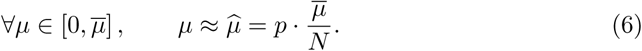

With this notation,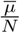 is, in fact, the resolution of the approximation. We can then perform the binary expansion of *p*:

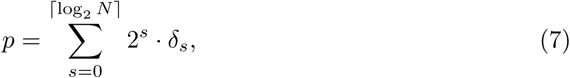

where ⌈log_2_ *N*⌉ denotes the smallest majoring integer to log_2_ *N*, and *δ*_*s*_ *∈* {0,1} is *r*^th^ digits from the right of the binary notation of *p*.

As an example, let us consider modeling an organism whose growth rate does not exceed *µ*_*max*_ = 2.3 h^−1^. To do this, we can set 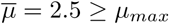. Let us choose a resolution of 0.25 h^−1^, which gives *N* = 10. Then, *log*_2_*N ≈* 3.32, and ⌈log_2_ *N* ⌉= 4. A growth rate *µ* = 1.4 will be approximated by:

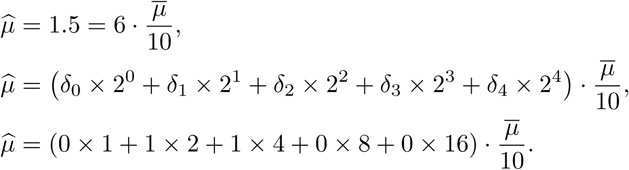

This example is illustrated in Fig. 1. Conceptually, a heuristic for solving this problem would be:

1. Solve the FBA for *µ*
2. Select the corresponding, closest 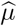
3. Apply it to compute dilution values
4. Solve the EP with fixed dilution
5. Apply the catalytic constraints to the FBA
6. Recalculate the FBA under catalytic constraints
7. If 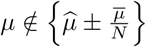, go back to 3, else, end.

**Fig 1.**
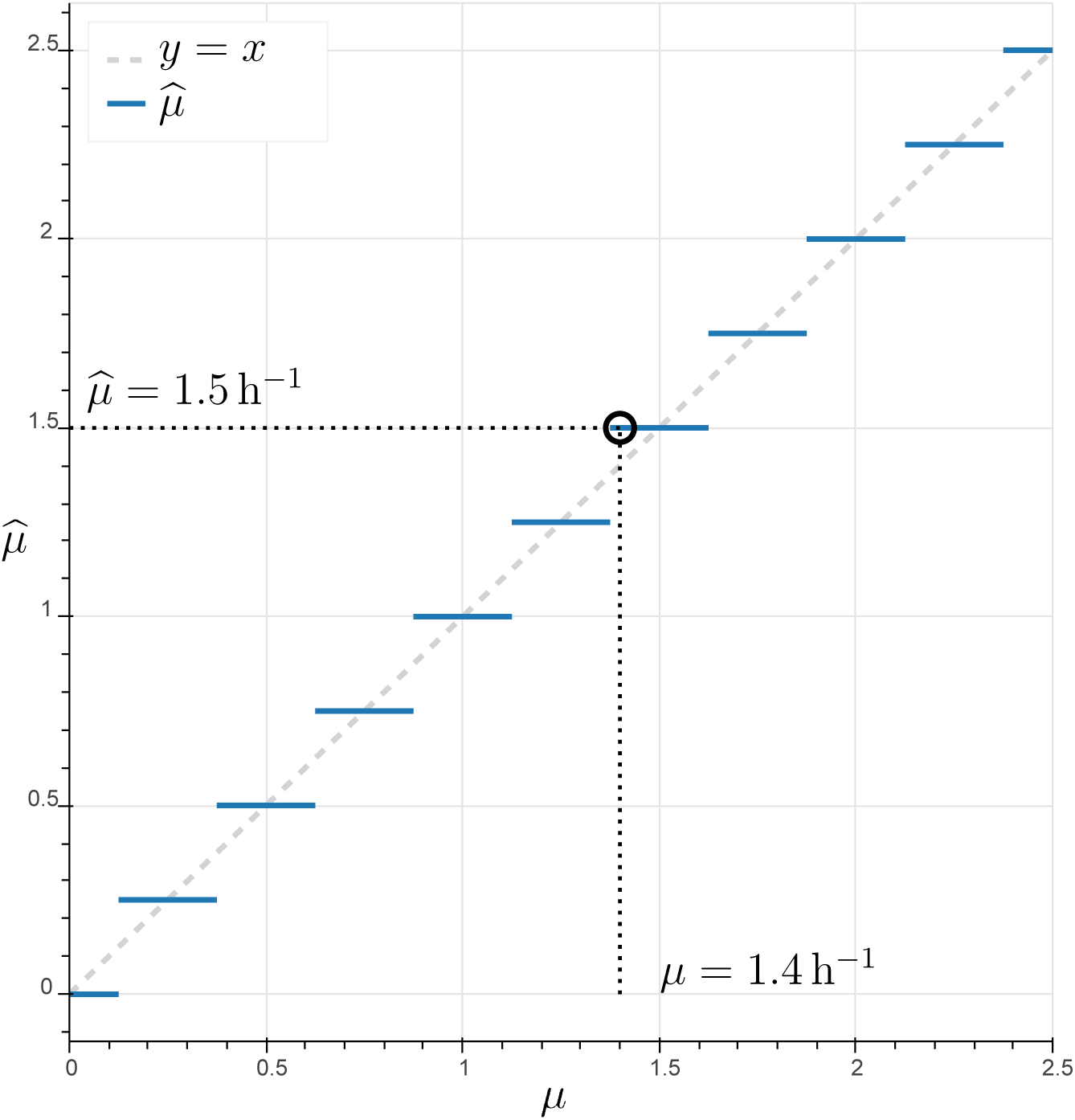
Discretization of *µ* into 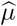. The step approximation transforms the continuous interval [0,1] into the discrete interval {0,0.25, …, 2.5}.

#### 1.2.3 Linearizing the bilinearity

In the previous derivation, we replaced the growth rate variable by a discrete number of acceptable values. We can approximate the continuous product *µ G*_⋆_, which represents the dilution, as follows:

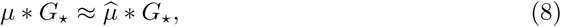

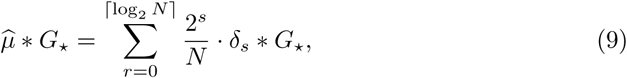

The product *δ*_*l*_ *∗ G*_⋆_ is then still bilinear, but one of its variables is binary. Assuming a constant *M > G*_⋆_, We can use Petersen’s linearization theorem [13, 14] to replace the product *δ*_*s*_ *∗ G*_⋆_ with a single nonnegative variable 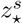, as described in the Materials and Methods.

Because of the binary expansion, the complexity of the model grows only as 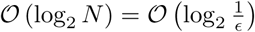, where ϵ = 1*/N* is proportional to the resolution of the approximation (which is 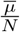). This means that the linearization part of a model with a resolution of 0.01 h^−1^ is only around twofold bigger than that of a model with a resolution 0.04 h^−1^, while resolution has been improved fourfold.

#### 1.2.4 Discretization of growth-dependent parameters

##### mRNA and enzyme content

Since growth has been discretized, it is now possible to also directly discretize other growth-dependent parameters of the problem, regardless of whether they are in a linear or non-linear relationship with growth. This is a direct consequence of the formulation of ETFL, which allows some flexibility in the modeling assumptions of the user. As an example, we described the relationship between growth and protein and mRNA mass ratios, *P*^*m*^ and *R*^*m*^, in the cell as reported in Neidhardt *et al.* [11] by interpolating and discretizing the protein ratio and mRNA ratio as functions of the growth rate. We used type 1 special ordered set constraints (SOS1) to model these variables with a first-order (piecewise linear) approximation:

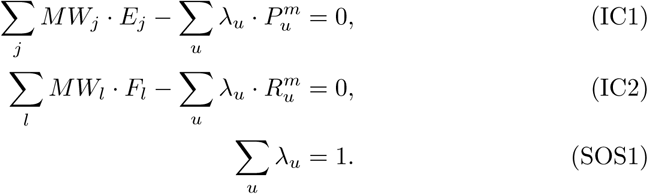

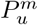 and 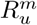 are growth-dependent, interpolated protein and RNA mass ratios (in g.g^−1^). Given a growth rate, they define the relative mass of the cell that is protein or RNA. *MW*_⋆_ represents the molar weight of the corresponding enzyme or RNA. The first two constraints enforce equality between the interpolated data and the model production. The last line is the SOS1 constraint that forces only one of the *λ*_*u*_ to be active.

Additionally, it is necessary to have the integer index of *λ*_*u*_ equal to the index of the growth rate. This is obtained through the constraint:

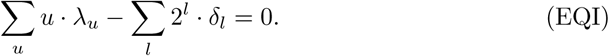

The first term represents the growth integer index, and the second represents its binary expansion. However, imposing such mass ratios requires the addition of a dummy mRNA as well as a dummy protein to represent the part of the transcriptome/proteome that is either missing from the expression model or altogether unrelated to metabolic function. We use average amino acid frequencies and GC content to model this. Explicit interpolation functions can also be used, such as the growth-dependent functions given in Pramanik *et al.* [17].

The simultaneous use of catalytic constraints on metabolic reactions (Eq. FC_j_, BC_j_) and maximal enzyme load (Eq. IC3) effectively implements allocation constraints like in GECKO [5], although in ETFL, the enzyme concentrations are also directly linked to the metabolism. In GECKO, the metabolic cost of building the enzymes is not taken into account.

Fig. 2 shows an example piecewise linear interpolation of the growth-dependent protein mass ratio in *E. coli* according to Neidhardt *et al*. [11]. The reported values (red circles) are interpolated using a piecewise linear function (dashed line), which is then discretized (full line). Using the integer constraints described above, the model can be forced to display a protein content that corresponds to its growth. We apply the same techniques to mRNA and DNA content.

**Fig 2.**
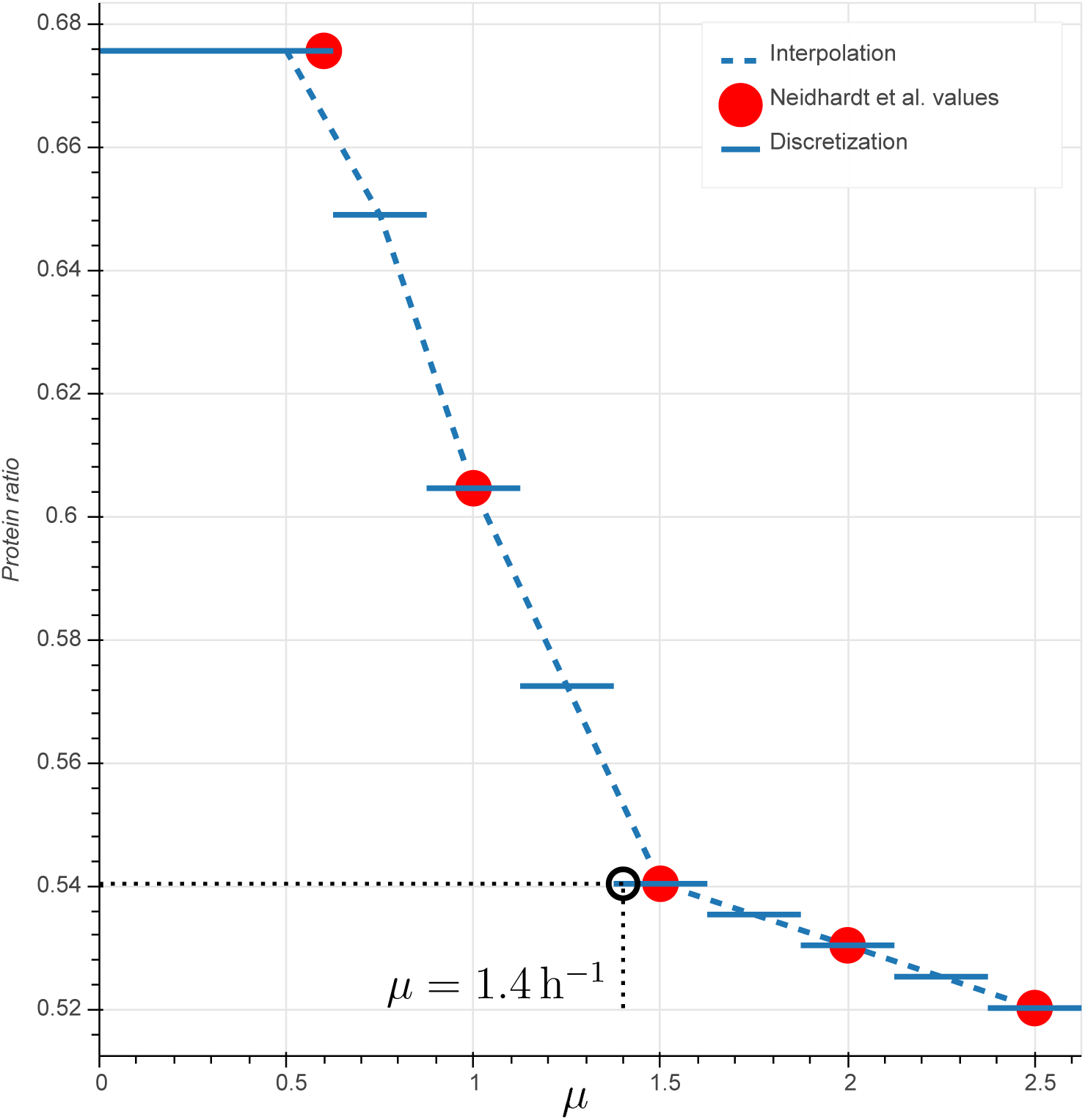
Example of piecewise linear interpolation and discretization of the protein mass ratio from Neidhardt *et al*. [11]. Red circles represent the values reported. The dashed line is the piecewise linear interpolation. The solid line is its discretization.

##### DNA content

To further increase the scope of macromolecules covered by the model, it is also possible to add growth-dependent DNA content. DNA mass ratios at specific growth rates are reported in Neidhardt *et al.* [11]. We model the DNA reaction synthesis as follows:

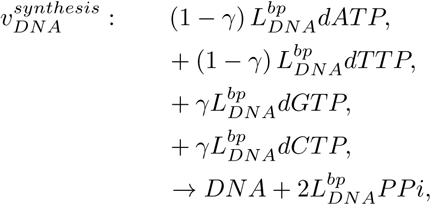

where *γ* is the GC content of the cell, and 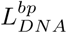 is the total length in base pairs of the DNA. As with *mRNA*_*l*_ and *Enz*_*j*_, *DNA* has a mass-balance equation of the following shape:

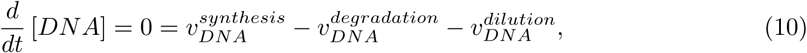

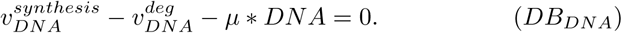

We consider that the DNA does not degrade, meaning the only source of DNA consumption is dilution caused by the growth of the cell and 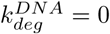. We then define the molar weight of DNA *MW*_*DNA*_ and enforce the DNA mass ratio *Dm* as we did with both proteins and mRNA:

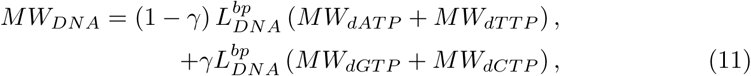

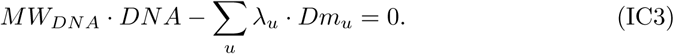

#### 1.2.5 Scale

A critical issue in the formulation of this problem is that the variables are different orders of magnitude. Fluxes are typically between 10^−3^−10^1^ mmol.gDW^−1^.h^−1^, whereas protein concentrations are around 10^−6^ −10^−3^ mmol.gDW^−1^ and mRNA concentrations are 10^−10^−10^−6^ mmol.gDW^−1^. The relationship between these scales is given by the catalytic rate constant of enzymes and expression machinery, which spans from 10^3^−10^6^*h*^−1^. In particular, the ribosome rate constant for translation (∼12 aa.s^−1^ = 43 200 aa.h^−1^) as well as the RNA polymerase rate constant of transcription (∼85 nt.s^−1^ = 306 000 nt.h^−1^) are responsible for strong differences in the concentrations and fluxes between transcription-and translation-related parts of the problem. Consequently, the constraint matrix becomes ill-conditioned, and the solver has to operate close to, or sometimes beyond, its maximal solving accuracy (usually around 10^−9^ for commercial solvers such as ILOG CPLEX or Gurobi).

To circumvent these limitations, we scale the EP, which will reduce the numerical difficulty of the problem, using nondimensionalization. We create nondimensionalized variables by dividing the variables of the initial problem by an estimated upper bound. For example, by definition, macromolecule concentrations cannot exceed 1 g.gDW^−1^, and the following constrains the transformed macromolecule variables between 0 and 1:

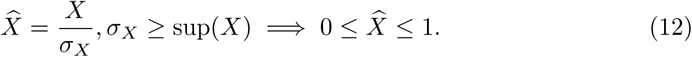

In this scheme, *σ*_*X*_ is an upper bound to *X*. In particular, if we consider *σ*_*X*_ to be the concentration of 1 g.gDW^−1^:

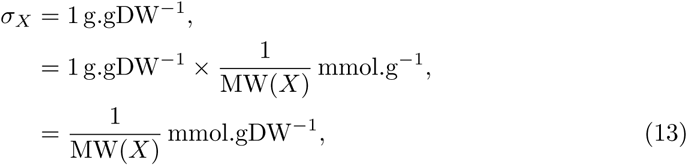

where MW(*X*) denotes the molecular weight of the macromolecule in SI units (kg.mol^−1^ ≡ g.mmol^−1^), and 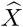 represents the mass fraction of the molecule in the cell. We scale the fluxes using a method derived from this, detailed in the supporting file S1 Nondimensionalization. It is also possible to further refine this upper bound by performing a variation analysis on *X* and re-generating a model using the newly estimated upper bound.

For the sake of clarity, all problem formulations will be kept in their dimensionalized form in the subsequent equations although the implementation is actually nondimensionalized. The nondimensionalized problem is described further in S1 Nondimensionalization.

#### 1.2.6 Recovering the FBA problem

In the ETFL formulation, enzyme synthesis is driven by the coupling between FBA and EP through the catalytic constraints. To carry flux, the cell needs to produce enzymes whose production will also use the metabolic resources of the cell. If allocation constraints are enforced, the amount of protein and mRNA synthesized must meet predefined mass ratios for the problem to be feasible. Hence, the metabolic requirement terms for the expression machinery (amino acids and NTP) have been removed from the biomass reaction and are accounted for in the tRNA charging and transcription reactions. Thus, the FBA solutions can be recovered from the ETFL formulation by the following routine:

- Setting 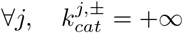,
- Constraining 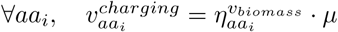,
- Constraining 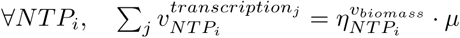,
- If applicable, relaxing the allocation constraints,
- If applicable, relaxing the thermodynamic coupling constraints.

### 1.3 Application: *E. coli* genome-scale model iJO1366

iJO366 [12] is a well-curated and well-studied GEM of *E. coli* that is closely related to the GEM used in developing both ME-models iOL1650-ME [7] and iJL1678b-ME [8]. Additionally, this model has been extensively applied in the literature and is aligned with a variety of datasets that can be used for data integration. We wanted to subject the model to classical studies that would highlight the power of ETFL, particularly as pertains to proteome-limited growth, macromolecule concentration variability analysis, and gene knock-out studies. We also wanted to assess the sensitivity of the model with respect to the presence of thermodynamic constraints as well as growth-dependent parameters.

Thus, we first experimented with four different models using ETFL with or without thermodynamic constraints and growth-dependent protein/RNA/DNA allocation following Table 2 as reported by Neidhardt *et al.* [11]. The following Table 4 details the nomenclature used to refer to these different models. The features of the most constrained model containing both thermodynamic and growth-dependent parameters, vETFL, are detailed in Table 5. These four models were optimized for maximal growth at increasing glucose uptake rates to assess their behavior with respect to excess substrate, which will show the non-linearity of the relationship between growth and glucose uptake at high uptake rates. A plateau in the growth rate was expected, which indicates a proteome-limited phenotype that cannot be observed with FBA. We also subsequently subject vETFL to a variability analysis and gene essentiality analysis, which will respectively show us the flexibility of the model and its accuracy in predicting gene knock-out behavior.

**Table 4.**
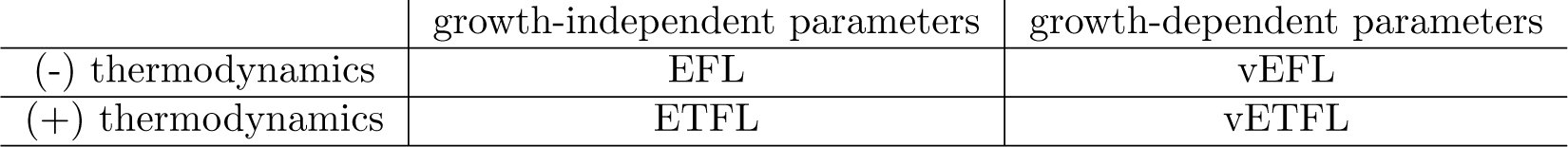
Nomenclature of the models used in the study of *E. coli* iJO1366. EFL stands for <u>E</u>xpression and <u>FL</u>uxes, ETFL for <u>E</u>xpression, <u>T</u>hermodynamics, and FLuxes, and the v-prefix indicates the inclusion of growth-dependent parameters (see 1.2.4)

|  | growth-independent parameters | growth-dependent parameters |
| --- | --- | --- |
| (-) thermodynamics | EFL | vEFL |
| (+) thermodynamics | ETFL | vETFL |

**Table 5.** Properties of the vETFL model generated from iJO1366.

|  |  |
| --- | --- |
| Growth upper bound $\hat{\mu}$ | $3.5h^{-1}$ |
| Number of bins $N$ | 128 |
| Resolution $\frac{\hat{\mu}}{N}$ | $0.0273h^{-1}$ |
| Number of constraints | 68304 |
| Number of variables | 49207 |
| Number of species | 3240 |
| – Metabolites | 1806 |
| – Peptides | 1431 |
| – rRNA | 3 |
| Number of enzymes | 562 |
| Number of reactions | 8023 |
| – Metabolic | 1543 |
| – Transport | 733 |
| – Exchange flux | 330 |
| – Transcription | 1431 |
| – Translation | 1431 |
| – Complexation | 562 |
| – Degradation | 1993 |
| Number of metabolites $\Delta_f G'^o$ | 1737 |
| Number of reactions $\Delta_r G'^o$ | 1787 |
| Percent of metabolites $\Delta_f G'^o$ | 93.9% |
| Percent of reactions $\Delta_r G'^o$ | 68.6% |

#### 1.3.1 Growth rate prediction

To study the behavior of the model at different carbon uptake rates, we simulated growth on a minimal medium with only glucose as a carbon source, unlimited oxygen, and some essential inorganic compounds. This would allow us to show that at a higher carbon uptake, the model would predict a limited growth – unlike FBA that would predict an unlimited linear increase.

Figure 3 shows the predicted growth rate of the different (v)E(T)FL models described in Table 4 with respect to the glucose uptake of the cell. As expected and in contrast to current FBA models, all four models plateau after a certain uptake rate, which indicates a proteome-limited phenotype due to the limited capacity of the cells to make more enzymes to metabolize the glucose. As discussed for the ME-models [7] and GECKO [5] formulations, within the context of models accounting for protein usage, this is caused by (i) the protein burden necessary to metabolize higher fluxes; (ii) the increased demand in protein synthesis at higher growth rates; and (iii) for the models with allocation constraints, the allowed protein and RNA mass ratio. We can see that models featuring protein, RNA, and DNA allocation constraints (vE[T]FL) consistently predict a lower growth rate than models without allocation constraints. This is expected, as the data we input requires additional proteins and mRNA to account for non-metabolism-related macromolecules. Models featuring thermodynamic constraints ([v]ETFL) also predict a lower growth rate, consistent with the fact that thermodynamic constrain the model to valid solutions whose flux is in the subspace of the FBA feasible space. The most constrained model (vETFL) consequently has the lowest growth rate at any glucose uptake. This is in accordance with published TFA results that eliminated biologically infeasible flux profiles yielding non-realistic higher growth rates [2].

**Fig 3.**
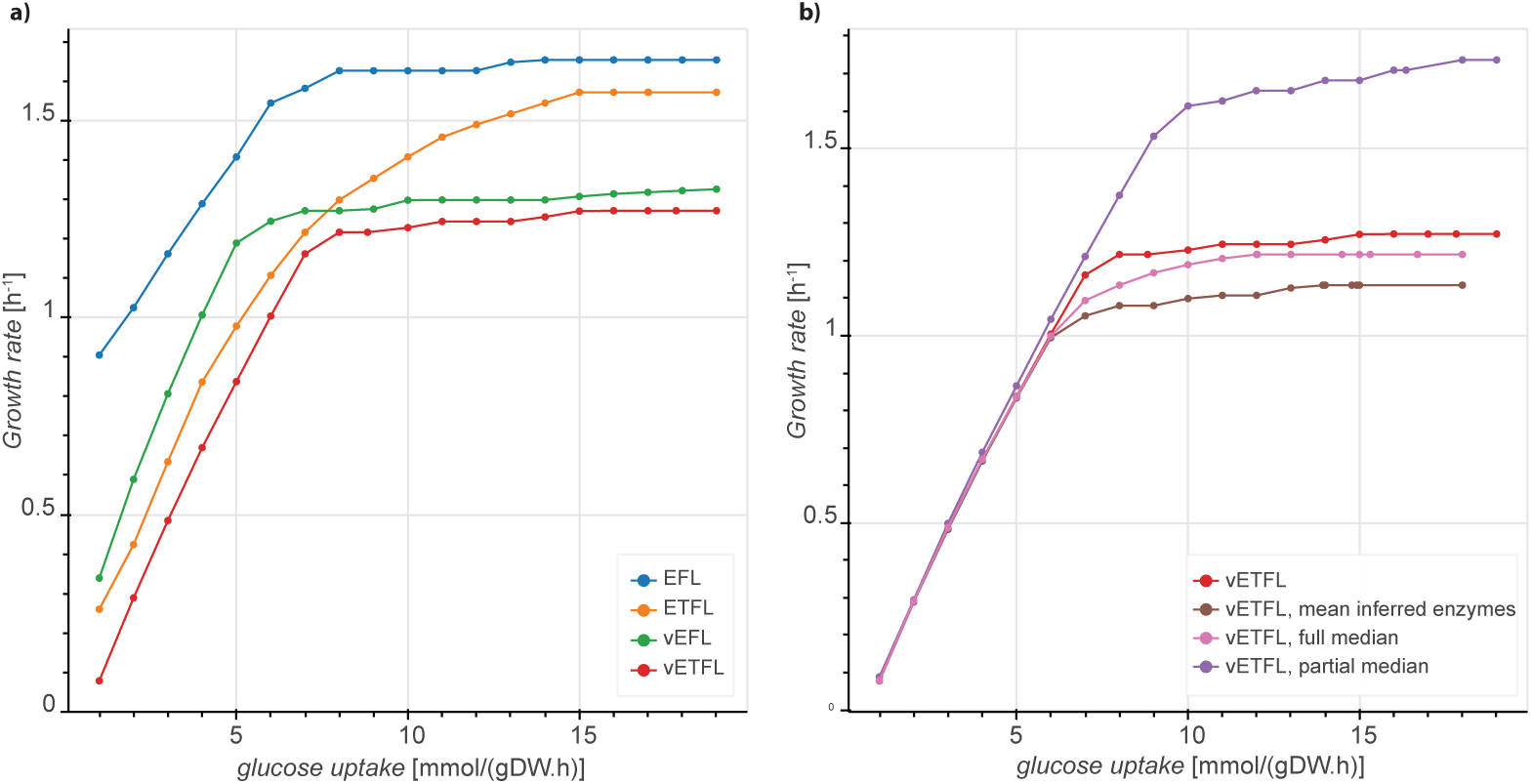
Growth rate with respect to glucose uptake for differently constrained models in the ETFL framework. **a.** Growth rate predictions using the [v]E[T]FL models with *k*_*cat*_ values aggregated from the reaction database SabioRK [18], **b.** Growth rate predictions accounting for missing enzymes using vETFL and models (i)-(iii) representing different initial enzyme assumptions, with *k*_*cat*_ values obtained from SabioRK or *k*_*cat*_ = 65 h^−1^, and with/without inferred enzymes.

We summarize the constraint matrix of the EP of vETFL in Fig. S2 Example EP constraint matrix, where each line represents a type of constraint and each column represents a type of variable. The blocks of the matrix that are non-zero are colored, and these blocks directly reflect the involvement of the constrained variables.

#### 1.3.2 Modeling missing enzymes

Although we initially focused on including only enzymes for which we had all the necessary information (catalytic rate and peptide constitution), we wanted to assess the behavior of our model when the missing enzymes were modeled as well as check our model’s sensitivity to changes in the catalytic rate constants. Thus, we additionally built three more models, based on vETFL, with the following properties: (i) all the missing enzymes were estimated by averaging the properties of the known enzymes based on the curation for the vETFL iJO1366 (333 amino acids long, average 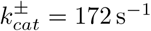); (ii) all the enzymes (including the missing enzymes) but the ribosome, RNA polymerase, and ATP synthase were assumed to have a catalytic rate constant 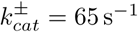; and, for comparison purposes, (iii) all the *known* enzymes of vETFL except for the ribosome, RNA polymerase, and ATP synthase were assumed to have a catalytic rate constant of 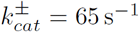. *k*_*cat*_ = 65 h^−1^ is an assumption used in O’Brien *et al.* [7] that we reproduce here as a study on sensitivity of the quality of the *k*_*cat*_ values. We observe that this value matches the median value of the catalytic rate constants in the 562 vETFL enzymes. For clarity, we will refer to these models as (i) the estimated mean enzymes model; (ii) the full median model; and (iii) the partial median model. The ribosome, RNA polymerase, and ATP synthase were not modified, as their catalytic rates directly and strongly affect the growth of the organism. Any drastic change in these would make changes related to other enzymes negligible in comparison.

Figure 3b shows a comparison of the growth prediction for the estimated mean enzymes (brown), full median (pink), and partial median (purple) models designed to account for the missing enzymes. For a better comparison, we also reproduce the vETFL results in red on the same graph. The partial median model (purple) shows a higher predicted growth than the original vETFL model (red). This implies that limiting enzymes in the original vETFL model have a *k*_*cat*_ parameter lower than the median value. Both models featuring inferred enzymes, the full median model (pink) and estimated mean enzymes model (brown) show a lower growth rate at a given uptake. This is expected as fluxes which previously had no enzymes assigned in vETFL are now subject to catalytic constraints, and thus the models are more constrained. Additionally, we observe that the model with estimated mean enzymes (brown) is also below the full median model (pink). Similarly to vETFL and the partial median model, this shows that the limiting enzymes in the model with estimated mean enzymes have a *k*_*cat*_ parameter lower than the median value. Finally, we observe that the differences between these four models only appear at glucose uptake rates higher than ≈6 mmol_glc_.DW^−1^.h^−1^, when the problem switches from being stoichiometry-limited to proteome-limited. Thus, this experiment also illustrates the importance of well-curated catalytic rate constants for modeling organisms grown in proteome-limited regimens.

These results demonstrate the capability of ETFL to predict different phenotypes depending on growth rate. ETFL is also amenable to hypothesis testing, as evidenced using the models that estimate the missing enzymes. In particular, we showed with ETFL that an uptake increase does not yield a proportional growth rate increase as with FBA and that ETFL provides a maximal uptake rate that is unmodeled in FBA, thus more effectively modeling growth-dependent biomass yield in *E. coli*. This allows for more realistic predictions for phenotypes that are limited by the expression capabilities of the cell as well as captures the variability of the biomass composition in different growth regimens.

#### 1.3.3. Variability analysis

It is also possible to subject the model to a range of variability analyses. These are routinely used in FBA to assess the flexibility of the system and in TFA to find the ranges of allowed metabolite concentrations. We can extend their use in ETFL to explore the allowed proteome and transcriptome. For example, we measured the admissible extreme concentrations of each mRNA in aerobic growth conditions as described in McCloskey *et al.* [19] by performing a variability analysis on the mRNA concentration variables. Fig. 4 depicts the admissible peptide concentration upper and lower bounds, sorted by average, for vETFL with a glucose uptake set to 12.5 mmol.gDW^−1^.h^−1^, which yields a proteome-limited phenotype, according to our results in Fig 3a. It is important to note that all peptides with a non-zero minimal concentration (most of the left of the figure) are, by definition, essential peptides: These are always present at this uptake rate and are hence necessary for the cell to grow at an optimum growth rate. The same study can be performed for enzyme concentrations or even metabolite log-concentrations for models with thermodynamics. This type of study is useful for comparing how the model performs in relation to actual proteomics, transcriptomics, or metabolomics data. The method for running these other types of variability analyses is exactly the same – only the variables subject to the variability analysis are changed.

**Fig 4.**
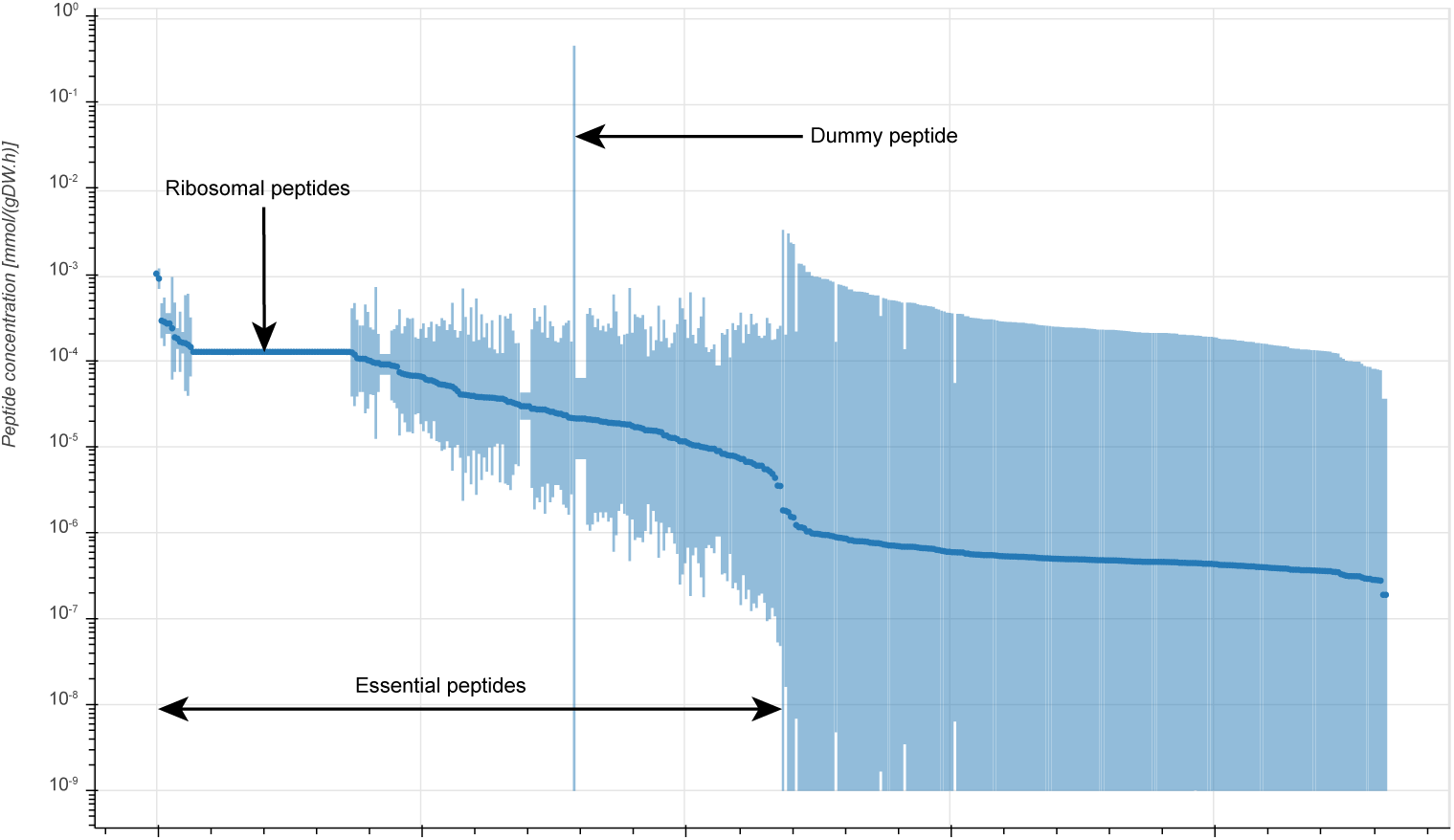
Concentration variability of peptide species, sorted by average peptide concentration (darker disc). Lower bounds that were 0 were set to the accuracy of the solver, 10^−9^. The horizontal line on the left side of the figure represents ribosomal peptides, which is narrow due to their instrumental role in making the tightly constrained amount of protein in the cell at a given growth rate. The vertical line in the middle represents the dummy peptide, which accounts for unmodeled peptides (non-metabolic proteins and enzymes with missing information) and therefore is used by the solver as a slack.

A specific usage of a variability analysis is the study of the allowed proteome (resp. transcriptome) that is done by performing a variability analysis on the enzyme (mRNA) concentration variables. This type of study can, for instance, be compared with transcriptomics to check if the expression profile of an engineered strain corresponds to what is expected in its corresponding model. A way to visualize the average allowed proteome (transcriptome) is to use the average value of the variability of each enzyme (mRNA) concentration as a feasible observation. Due to the convexity of the solution space, it is a solution to the problem. This observation is then plotted on a finite area, which can be done using the online software Proteomaps [20, 21]. This method and software are often used by biologists to represent protein abundances in the cell, and using the data from ETFL, we can generate similar comparative graphs that can help biologists analyze the variability in the different concentration variables using a visualization they are familiar with.

Fig. 5 is an example of such a representation, graphed using the mRNA concentrations corresponding to the solution represented by the dark dots in Fig 4 as an input. In this figure, mRNAs are clustered using KEGG Gene Ontology (GO) annotations. GO annotations form a tree describing the physiological role of genes, ranging from the least specific (e.g. general metabolism) to most specific (e.g. araH gene). The area of each (sub)cluster is proportional to the relative abundance of each (sub)group of mRNAs.

**Fig 5.**
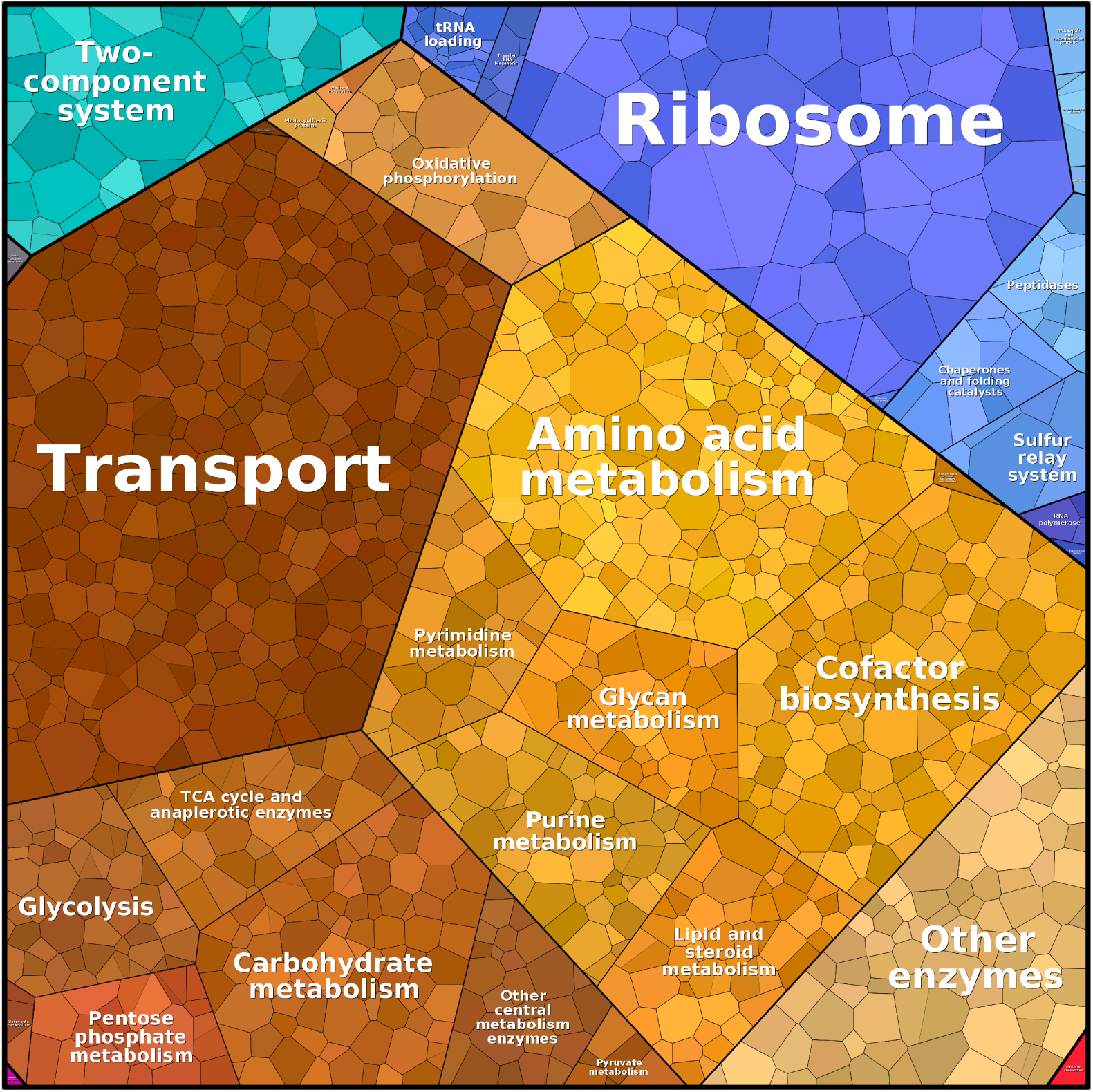
Proteomaps of the predicted abundances of mRNAs. Each colored patch represents a different mRNA in proportion to its relative abundance. Genes can be clustered using KEGG Gene Ontology (GO) terms.

We used the mean of the variability analysis as the observation rather than a single optimal solution because the optimality principle in LP only guarantees a unique global optimum value and not a unique optimal solution. Moreover, solver heuristics give sparse and extreme results (corners of the explored simplex), which do not accurately represent the full extent of the considered solution space.

#### 1.3.4 Essentiality analysis

The ETFL framework can also analyze the essentiality of specific genes by performing single gene knockouts. The growth of models with knocked-out genes can then be compared to experimental data to assess the quality of the model as a validation. We performed this analysis and compared it to the results reported in the publication of iJO1366 by Orth *et al.* [12]. We use the Matthew’s correlation coefficient (MCC) as a metric for the quality of the prediction, which is preferred over accuracy as it is not sensitive to the imbalance between the number of essential genes and non-essential genes. The MCC reads like a usual correlation coefficient, with 1 being a perfect correlation, −1 perfect anti-correlation, and 0 no correlation. We used the essentiality data and conventions given in the supplementary material of Orth *et al.* [12], as explained in Fig. 6-a and Fig. 6-b. Knockouts in ETFL are simulated by setting the corresponding transcription reaction rate to 0 as opposed to in FBA where knockouts derive from Gene Protein association Rules (GPR), which are Boolean rules that directly set metabolic reaction rates to 0 if the knocked-out gene is essential to this reaction. The difference is that in ETFL, the solver actually sees the knock-out as a part of the model, while FBA changes the model to accommodate the knock-out. This feature can be used in strain design strategies to optimize directly for knock-outs. The results are presented in Fig. 6-c and 6-d, respectively for vETFL and the partial median model.

**Fig 6.**
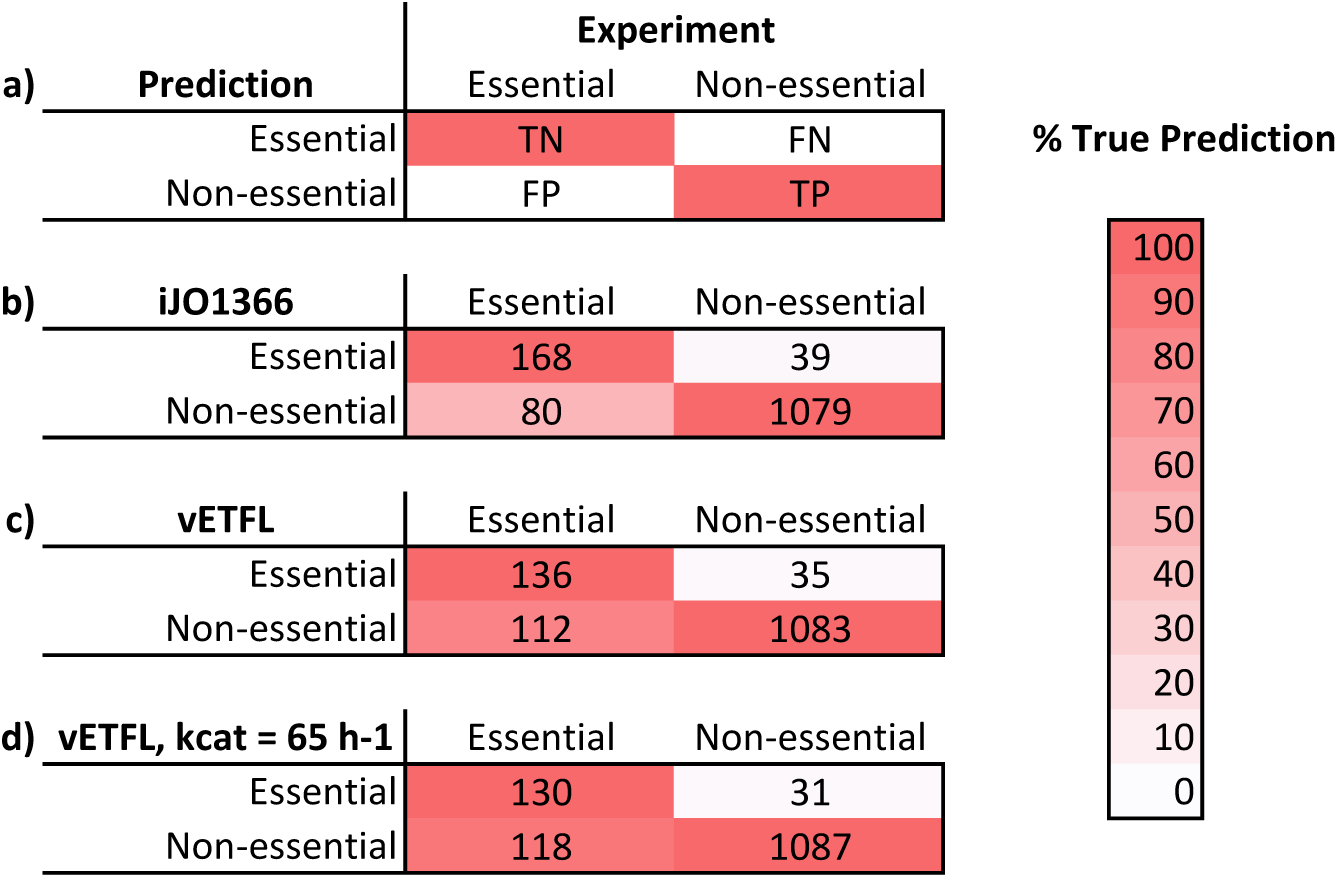
**a.** Conventions from Orth *et al.* [12] for gene essentiality. TN is True Negative. FN is False Negative. FP is False Positive. TP is True Positive. The color shading represents how good the classification is in the experimental class. Perfect classification should have a strict red first diagonal, as shown on this example. **b.** Gene essentiality prediction for FBA model iJO1366, yielding a Matthew’s correlation coefficient (MCC) of 0.69. **c.** Gene essentiality prediction for vETFL model, yielding a MCC of 0.60. **d.** Gene essentiality prediction for vETFL model with all *k*_*cat*_ = 65 h^−1^, yielding a MCC of 0.59

We observe that, compared to iJO1366, (i) ETFL presents more false positives (experimentally essential genes predicted as non-essential); and (ii) ETFL predicts fewer false negatives (experimentally non-essential genes predicted as essential). This first point might indicate that ETFL is less constrained than iJO1366—the cell has more genetic alternatives for growth. This is an artificial effect that likely stems from the missing enzyme data, because if a reaction depends on a missing enzyme, by default there will be no coupled expression and hence no impact if the related gene is knocked out. As more enzyme data is added to the model, the false positive rate should decrease. The second point indicates that ETFL better captures the genes that are not essential for growth than iJO1366. This is an expected effect of explicit expression coupling, from transcription to enzyme concentration, as opposed to Boolean gene reaction rules.

#### 1.3.5 Sampling

Sampling the feasible solution space of FBA is a common way to study solution robustness and variability. Since there are often multiple FBA solutions at the optimal objective value, representative solutions are often sought, and sampling is one way to obtain them. However, because ETFL contains integer variables, it is not compatible with traditional sampling methods in its current formulation. It is possible, though, to make the model convex, and hence amenable to sampling, by fixing the integers to their values at a given growth rate and, if applicable, TFA directionality. This will block the flux directions as well as the growth-dependent parameters if TFA is performed. The resulting model is then solely linear, and sampling can be performed with traditional techniques, such as artificially centered hit and run (ACHR) [22], gpSampler [23], or optGpSampler [24]. Once it has converged, a sampling should provide a better representation of the center of the solution space than the mean of the variability analysis.

### 1.4 Performance

ETFL relies on solver-specific MILP algorithms and heuristics, which also means that great variability in performances can be observed depending on the solver parameters. We provide tuned presets for different tasks (gene knock-out, variability analysis, growth maximization) with the package, and recommend that users run their own solver tuning if long run times are observed. We witnessed an up to 10× increase in performance using such tuning.

### 1.5 Adaptation of FBA-based methods to ETFL

The ETFL formulation is amenable to further kinds of analyses. Leveraging both the explicit expression constraints and the MILP nature of the problem, we present several possibilities for future studies using ETFL:

#### Growth-dependent parameters

It has been reported that several other parameters, such as the ribosome transcription rate constant *k*_*rib*_, are growth dependent [11]. Although such dependency is not taken into account in the presented results, it is possible to account for this by (i) discretizing *k*_*rib*_ following the method used in 1.2.4, and (ii) using Petersen’s linearization scheme from 1.2.3 on the product *k*_*rib*_ ∗ *E*_*rib*_. Other parameters that can be transformed in this way include, but are not limited to, the RNAP transcription rate constant *k*_*trans*_, free ribosomes, and the RNAP ratios *ρ* and *π*.

#### Omics integration

Explicit mRNA and enzyme concentrations allow the direct integration of absolute or relative proteomics and transcriptomics by changing the bounds of the corresponding variables in the EP. An additional gauge constraint will be needed for relative data. Previous transcriptomic integration methods, such as REMI [25], iMAT [26], GIMME [27], or MINEA [28], can also be adequately reformulated for ETFL. Metabolomics can still be integrated using TFA [2, 3].

#### Minimization of adjustment

In the original paper, the hypothesis behind the Minimization of Metabolic Adjustment (MOMA) method is that the metabolic fluxes of an organism subject to a gene knock out show a minimal change compared to the metabolic fluxes of the wild-type organism [29]. The underlying hypothesis is that the enzyme distribution and assignments remain the same except for the knocked-out gene. With ETFL, it is possible to directly compute a Minimization of Protein Adjustment (MOPA) by reformulating the objective function as a Minimization of Expression Adjustment (MOXA):

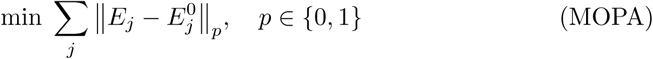

where ‖ *·*‖_*p*_ is either the Manhattan norm (*p* = 1, *ℓ*_1_-norm) or the Euclidean norm (*p* = 2, *ℓ*_2_-norm), which will require a MIQP solver. In the same fashion, it is also possible to formulate a (weighted) Minimization of mRNA Adjustment (MORA) or even a Minimization of eXpression Adjustment (MOXA) using the following formulations:

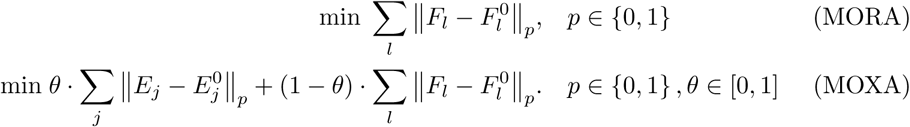

#### Parsimonious analysis

Parsimonious FBA (pFBA) [23] was developed to address the high fluxes of some of the solutions given by FBA. Although this concern is addressed in ETFL by the combined actions of the EP and thermodynamics, pFBA can be adapted to ETFL to study an organism under parsimonious constraints. For example, it is possible to reformulate it into a parsimonious expression problem to find the minimal expression level required to meet a growth target using objective functions similar to MOPA, MORA, and MOXA. It is also possible to turn the problem around to consider the allowed enzyme amounts under minimal flux constraint obtained by pFBA to assess the metabolic flexibility of an organism.

#### Dynamic ETFL (dETFL)

Dynamic FBA (dFBA) [30] is a method that uses FBA to predict the dynamics of a biological system represented with a stoichiometric model. In its original static optimization approach (SOA) formulation, a FBA problem is solved at each time step. The value of boundary fluxes of the FBA problem are updated at each iteration with values produced with a kinetic law, such as Michaelis-Menten glucose uptake and oxygen diffusion. Because ETFL allows direct access to enzyme concentrations, it is possible to use the latter to reformulate dFBA in its SOA. The original SOA approach uses ad-hoc constraints on the absolute flux change at each time step. However, in ETFL, it is possible to bound flux changes indirectly by bounding enzyme and mRNA concentration changes in the EP. Effectively, this approach allows the movement from ad-hoc constraints to physiological constraints.

#### Use in kinetic frameworks

Often, kinetic frameworks require a reference flux distribution as an input. ETFL can provide such a distribution, with an increased accuracy as compared to FBA.

## 2 Conclusions

ETFL is a framework can implement expression and thermodynamic formalism using mainstream MILP solvers. This could not be previously accomplished using state-of-the-art ME-models, which use specialized quad-precision solvers and do not support integer variables. The formalism itself is based on the explicit and direct relationship with the underlying biochemistry and provides a way to incorporate growth-dependent variables using MILP linearization techniques. These new growth-dependent variables provide a finer modeling of expression because they consider phenotypic differences in different growth regimens, which is key for accurate modeling.

ETFL can also compute explicit mRNA and enzyme concentrations as well as perform directomics data integration. In this, ETFL complements and extends FBA capabilities by using explicit relationships in lieu of the typical assumptions on the relationships between the transcriptome, proteome, and fluxome. This explicit accounting of expression mechanisms provides a finer level of control and a more relevant prediction of gene-editing outcomes. Because of this and its operational similarity with classic FBA-related analyses, ETFL can be efficiently integrated in standard model-based pipelines. For example, metagenome-based genome-scale reconstructions such as published by Magnúsdóttir *et al.* [1] can be directly fed to the framework to generate one models for each of the 773 bacteria they identified. In a more general way, ETFL can assess the allowed expression profiles of any biological system amenable to genome-scale modeling, such as the metabolic engineering of biocatalysts, microbial communities, drug design, or personalized medicine.

## 3 Materials and Methods

### 3.1 Formulation of the expression problem

Several parameter values were taken from the BioNumbers database [31], and when used, their identification number as well as the original source from which the value was reported are specified.

#### 3.1.1 Preliminaries, Conventions, and Notations

We will write the mass balances for the macromolecules with included concentration variables. Because we assume the cell is growing at a growth rate *µ*, we must assume that the volume in which the mass balance is calculated varies.

The time derivative of the concentration of a macromolecule *G* of concentration *C*_*G*_ in the volume *V*, for a total mass *m*_*G*_ in the cell, produced at a rate 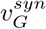 and degraded at a rate 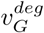, will be written:

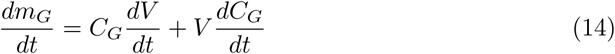

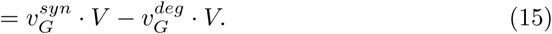

We can combine equations 14 and 15 and divide by *V* (necessarily non-zero) to write the time derivative of the concentration *C*_*G*_:

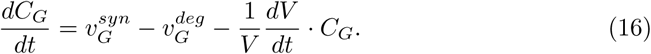

By definition, 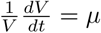 is the growth rate of the cell, and we the term *μ*. *C*_*G*_ the dilution term, or 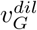, as per Fredrickson’s work on formulating growth models [32]. It is a common assumption that the concentrations inside the cell remain time invariant (quasi-steady state assumption), effectively yielding the constraint:

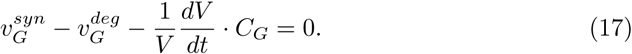

It is also understood from the formulation of the FBA that adding a new reaction to the system, such as:

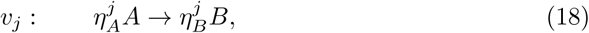

results in adding terms to the mass balances of *A* and *B*:

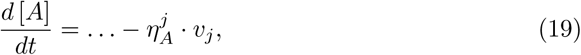

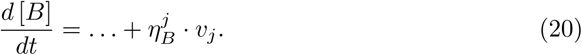

The further extension of this to reactions of n reactants to m products is trivial.

Several parameter values are taken from the BioNumbers database [31]. When used, we specify their identification number as well as the original source from which the value was reported. Finally, we will represent products between a parameter value and a variable by the symbol “·” and products between two variables by the symbol “*”.

Hereafter, we propose a detailed top-down approach to formulate the constraints being built for ETFL, starting from the metabolite network and moving down to RNA synthesis.

#### 3.1.2 Metabolic reactions

From FBA, the mass-balance relationship for metabolites can be written as:

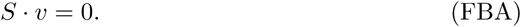

For the rest of the formulation, it is necessary to split the net flux *v* from each reaction into its forward net component and backward net component:

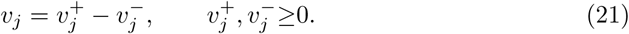

Biochemical reactions are catalyzed by enzymes. Each enzyme (*Enz*_*j*_) of concentration *E*_*j*_ can catalyze a flux *v*_*j*_ subject to the enzyme capacity constraint, which is a function of its forward and backward catalytic rate constants 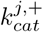 and 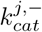:

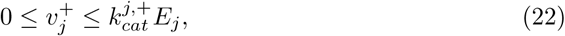

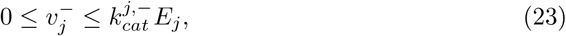

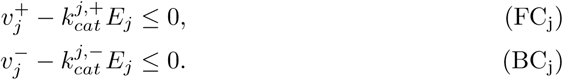

The distinction between the bounds of the forward and backward net fluxes is important, as some enzymes have different catalytic activities depending on the direction of the flux.

#### 3.1.3 Enzyme assembly

Each enzyme *Enz*_*j*_ in concentration *E*_*j*_ is subject to mass balance, which can be written:

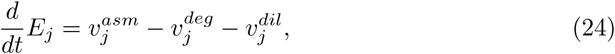

which reads under quasi-steady state assumption (QSSA):

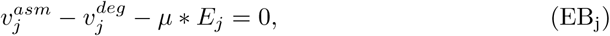

where 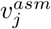 is the formation rate of the enzyme by the assembly of its constituent peptides, 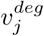 is the degradation rate, 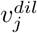 is the dilution rate, and *µ* is the growth rate of the cell. The formation rate of the enzyme describes the assembly of free peptides, so it is necessary to add the peptide assembly reaction to the stoichiometric matrix:

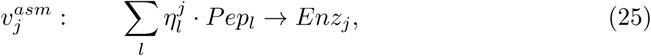

where 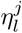 is the stoichiometric coefficient of peptide *Pep*_*l*_ for the formation of the complex of enzyme *Enz*_*j*_.

#### 3.1.4 Peptide synthesis

The synthesis of peptides consumes charged tRNAs, which are subsequently uncharged during the current peptide synthesis by a ribosome. The process consumes 2 GTP and releases 2 GDP and 2 Pi per amino acid:

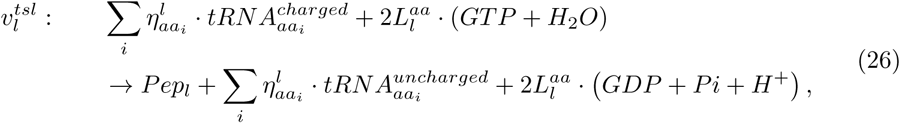

where *aa*_*i*_ denotes the *i*^th^ amino acid, 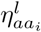 its relative stoichiometric coefficient (count) in the sequence of *Pep*_*l*_ 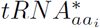 the (un)charged tRNAs for each amino acid, and 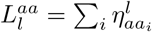 is the length of the amino acid sequence of *Pep*_*l*_.

As explained in section 1.1, this reaction adds a supplementary term in the mass balances of the metabolites (GTP, GDP, Pi, H_2_O, H^+^), the peptide, and the tRNAs (see 3.1.9 for the latter). This term is what connects the expression requirements to the metabolic network defined in the FBA.

In particular, the peptide concentrations obey the mass-balance equation:

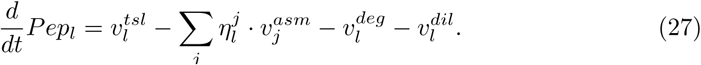

We assume in the current model that the assembly rates are much faster than dilution and degradation, and thus simplify this mass balance to:

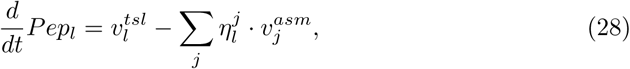

which, under QSSA, can be written:

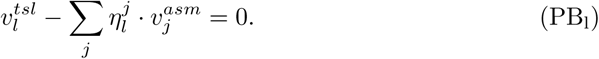

In this context, the peptides are treated just like regular metabolites in the system. This assumption in PB_l_ can be relaxed without a loss of generality by introducing a dilution and a degradation term, thus introducing a bilinearity.

Finally, we must take into account the actual degradation reactions. We add the ideal degradation reaction to the system:

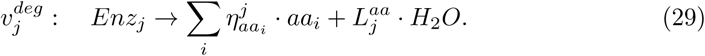

For this, the degradation rate is known:

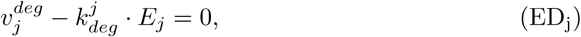

where 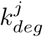 is the degradation rate constant of the enzyme.

#### 3.1.5 Ribosome synthesis and utilization

The peptides are the product of a translation reaction that is catalyzed by a ribosome. As we did with the catalytic constraints for general biochemistry reactions, we can apply the ribosome maximum catalytic rate as an upper bound to its translation rate 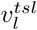:

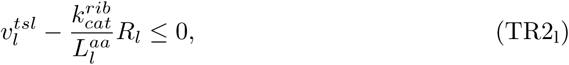

where 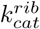 is the maximum ribosomal translation rate constant (10 - 12 aa.s^−1^ for *E. coli*, BioNumbers ID [BNID] 100059 [33]), 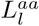 is the amino acid length of the peptide *l*, and *R*_*l*_ is the quantity of ribosomes assigned to the translation of this peptide. This way, the ratio *R*_*l*_*/Pep*_*l*_ is effectively the average polysome size translating the peptide *l*.

Like any other enzyme, ribosomes verify the mass balance:

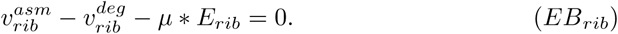

*E*_*rib*_ denotes the total quantity of ribosomes in a cell. It accounts for *R*_*l*_, the ribosomes assigned to the translation of *Pep*_*l*_, as well as the free ribosomes in the cell, *R*_*F*_. We can then write the total ribosome capacity constraint:

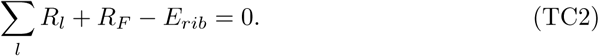

If we know the ratio *ρ* of free versus occupied ribosomes, we can enforce it:

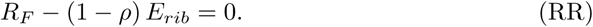

Finally, the ribosome differs from other enzymes in that it takes ribosomal peptides *rPep*_*l*_ as well as ribosomal RNA *rRNA*_*l*_ for its assembly. Hence, its assembly reaction is:

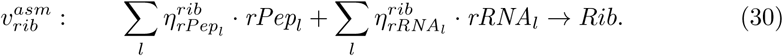

As explained earlier, the stoichiometric coefficients 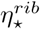 will appear in the mass balances of each of the compounds of the reaction.

#### 3.1.6 RNA Polymerase

The transcription is catalyzed by RNA polymerase (RNAP). For each transcription of mRNA, we can put an upper bound on the transcription rate 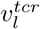 in the same way as for translation:

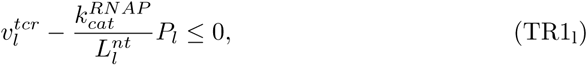

where 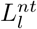 is the length in nucleotides of the mRNA sequence, 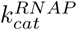 is the catalytic rate constant of RNAP (85 nt.s^−1^ for *E. coli*, BNID 100060 [33]), and *P*_*l*_ the concentration of RNAP assigned to the transcription of this mRNA.

RNAP is an enzyme, and hence it also satisfies mass balance:

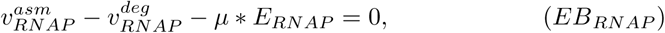

where *E*_*RNAP*_ is the total amount of RNAP, which also accounts for free RNAP *P*_*F*_, and follows:

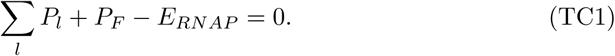

As we did with the ribosomes, if we know the ratio of occupied RNAP, *π*, we can enforce it:

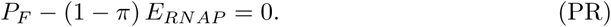

#### 3.1.7 mRNA synthesis and mass balance

During the translation, an mRNA is read to produce a peptide. mRNAs are subject the same mass-balance constraints:

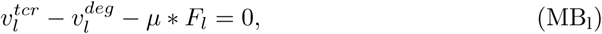

where *F*_*l*_ is the total concentration of the *l*^th^ mRNA (*mRNA*_*l*_), 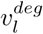 is its degradation rate, and 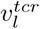 is its transcription (synthesis) rate. *F*_*l*_ is to *mRNA*_*l*_ what *E*_*j*_ is to *Enz*_*j*_ –the concentration variable that represents the macromolecule in the EP. The transcription reaction is modeled as follows:

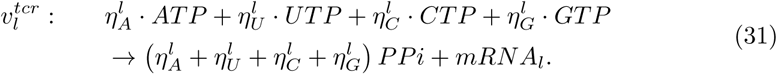

Again, the stoichiometric coefficients will appear in the mass balances of each of the metabolites and macromolecules involved.

We must also take into account the relationship between ribosome assignment and mRNA concentration. On each strand of *mRNA*_*l*_, there can be only a finite number *ρ*_*l*_ of ribosomes translating at the same time. This number is given by the ratio of the footprint size of the ribosome 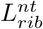 and the length of the mRNA strand 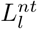. This effectively yields the number of ribosomes that can be present at the same time on a given mRNA strand:

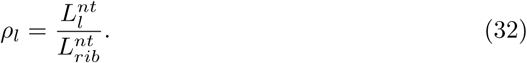

For *E. coli*, 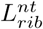 is approximately 20 nm (BNID 102320 [34], 100121 [35]), which amounts to approximately 60 base pairs (the length of a nucleotide is approximately 0.3 nm; BNID 103777 [36]). From there we can get the additional constraint:

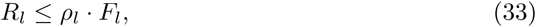

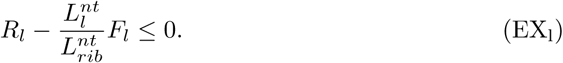

Finally, we must take into account the actual degradation reactions. We consider perfect degradation for mRNAs:

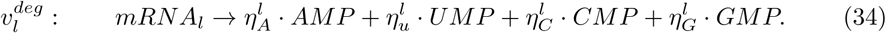

And, again, we know the degradation rates:

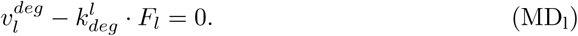

#### 3.1.8 rRNA synthesis and mass balance

*r*RNAs are used in the ribosome assembly reaction. According to the definition of 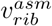 in 3.1.5, their mass balance can be written:

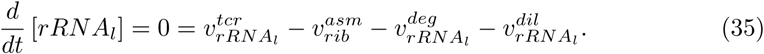

We neglect their dilution and degradation under the hypothesis that free rRNAs are scarce and stable [37]. Thus, their mass balance in the model reads:

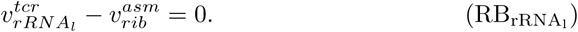

#### 3.1.9 tRNA synthesis and mass balance

tRNAs are produced with a charging reaction and consumed by peptide synthesis. We use the following charging reaction:

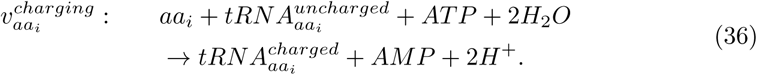

Once again, the stoichiometric coefficients of each reactant will appear in the stoichiometric matrix in the column corresponding to this reaction.

Since tRNAs are relatively stable molecules [37], we neglect their degradation. Let 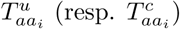 represent 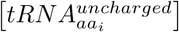 (resp. 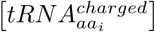). Then, we can write the following constraints:

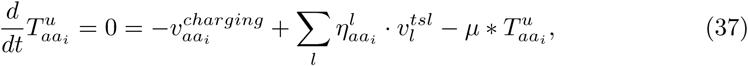

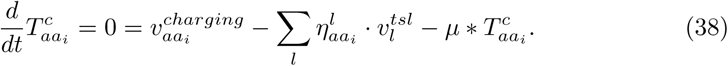

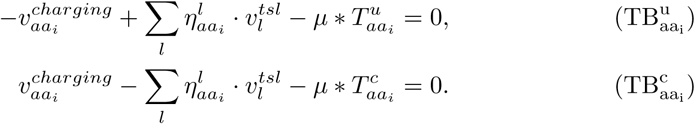

### 3.2 Thermodynamics-based constraints

Thermodynamics flux analysis (TFA) [2, 3] imposes constraints on a FBA problem to couple reaction directionality to the standard free energy of reactions and metabolite concentrations. We also introduce constraints that couple the sign of the Gibbs energy of a reaction to its directionality through the use of integer variables and a mixed-integer linear coupling formulation. This framework reduces the feasible flux space and improves the predictive power of FBA by removing thermodynamically invalid flux profiles.

Considering *c*_*i*_ is the concentration of *i*^th^ metabolite, we define *C*_*i*_ as its scaled logarithm with respect to *c*_0_ so that in standard conditions *c*_0_ = 1 M:

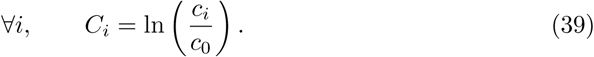

We use the 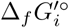 to calculate the 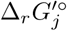 the standard Gibbs energy of formation in solution of the *i*^th^ metabolite and Gibbs energy in solution of the *j*^th^ reaction, respectively, and we obtain the additional variables:

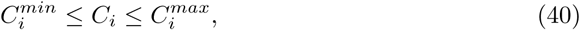

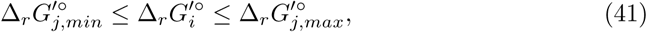

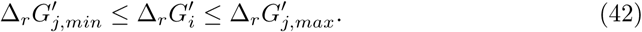

The concentration variables are bounded by experimental measurements or physiological assumptions, and the standard Gibbs energies are bounded by the measurement or estimation error. Since the net flux of each reaction has already been split between forward flux 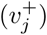 and backward flux 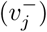, (see Eq. 21), we can directly add the constraints described in [2]:

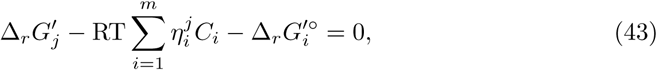

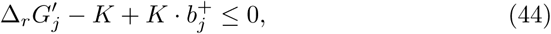

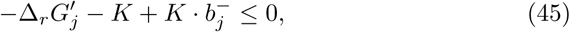

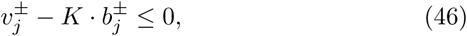

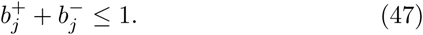

R denotes the ideal gas constant, T is the temperature in Kelvin, and 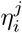 represents the stoichiometry of the metabolite *i* in the reaction *j*. *K* is a big-M constant (bigger than all upper bounds), and 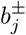 are binary variables. Eq. 43 defines the actual Gibbs energy of the reaction as a function of its standard Gibbs energy and the scaled logarithms of metabolite concentrations. Eq. 44 and Eq. 45 ensure that 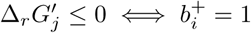 and 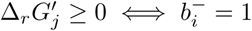. These binary variables are used to block flux in Eq. 46 if the thermodynamics do not favor it. Finally, Eq. 47 is added to enforce that only one direction is chosen.

### 3.3 Petersen linearization

After discretization of the growth rate, the dilution term for the macromolecule *G*_⋆_ will consist of a sum of products of the binary variables *δ*_*s*_ and the continuous variable *G*_⋆_We can use the Petersen linearization scheme [13] to transform this product into an equivalent system of one new variable and three new constraints:

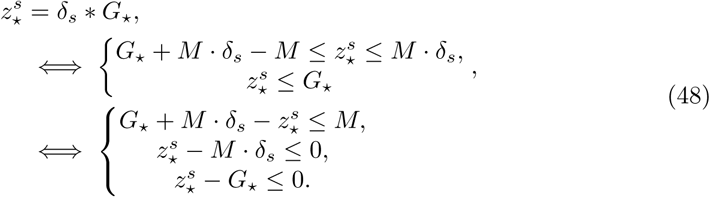

With this method, we can directly reformulate generalized mass balances as described in Eq. 5 for mRNAs, enzymes, uncharged tRNAs, and charged tRNAs:

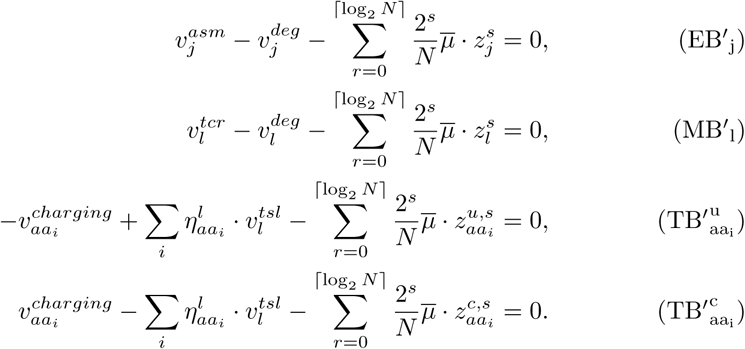

And we get the additional linearization constraints:

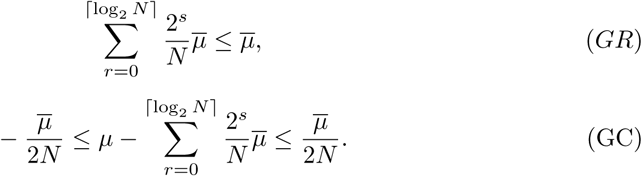

### 3.4 Data

#### 3.4.1 Degradation rate constants *k*_*deg*_ (proteins,mRNA)

mRNA degradation rates were taken from Bernstein *et al*. [38]. We converted the reported half lives into rate constants using the classical relationship 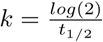. Enzyme degradation rates were approximated to 5% per hour, as per a report indicating this rate was between 2% and 7% per hour in [39].

#### 3.4.2 Catalytic rate constants *k*_*cat*_ (homomers,other complexes)

Catalytic rate constants 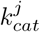 were obtained from Davidi *et al*. [40] for homomer enzymes. Complex formation reactions for non-homomer enzymes were taken from the supplementary information of O’Brien *et al.* [7] and Lloyd *et al*. [8]. EC numbers were obtained from BiGG [41] and the iJO1366 publication [12]. Their corresponding *k*_*cat*_ values were assigned using conservative (max) values from SabioRK.

#### 3.4.3 Peptide composition of enzymes

Homomer compositions were obtained from Davidi *et al*. [40]. Other peptide compositions of enzymes were taken from the supplementary information of O’Brien *et al*. [7] and Lloyd *et al*. [8]. Additional information was obtained from the Metacyc/Biocyc database [42, 43] using specialized SmartTables queries [44].

### 3.5 Modeling

#### 3.5.1 Model modifications

The initial model was subjected to minor changes to accommodate for ETFL modeling. In particular, we added:

- Selenocysteine as a metabolite.
- Cysteine to selenocysteine conversion as a pseudo reaction.
- tRNA charging reactions.

We also modified the biomass reaction by removing its nucleotide and amino acid components, since they are already taken into account by the expression problem as explained in 1.1.1.

#### 3.5.2 Enzyme estimation

Given a reaction in the model, if no enzyme is supplied but the reaction possesses a gene reaction rule, it is possible to infer an enzyme from it. The rule expression is expanded, and each term separated by an `OR` boolean operator is interpreted as an isozyme, while terms separated by an `AND` boolean operator are interpreted as unit peptide stoichiometric requirements. The enzyme is then assigned an average catalytic rate constant and degradation rate constant.

#### 3.5.3 Essentiality analysis

For increased performance, the essentiality analysis was cast into a feasibility problem. We put a lower bound on growth equal to 10% of the predicted ETFL growth and set the objective to 0. With this method, essential genes will cause the problem to be infeasible, while non-essential genes will return a feasible solution satisfying at least 10% of the growth. This method achieved up to a 5-fold reduction in solving time on the most complex models.

### 3.6 Computation

#### 3.6.1 Implementation

The code has been implemented as a plug-in to pyTFA [45], a Python implementation of the TFA method. It uses COBRApy [46] and Optlang [47] as a backend to ensure compatibility with several open-source (GLPK, scipy, …) as well as commercial (CPLEX, Gurobi, …) solvers. The code is freely available under the APACHE 2.0 license at https://github.com/EPFL-LCSB/etfl. We rely on the Python package Biopython [48] for transcribing and translating sequences of nucleotides and amino acids.

#### 3.6.2 Hardware

Computations were done on a 64-bit Ubuntu 18.04.1 LTS (Bionic Beaver); 2 ×Intel(R) Xeon(R) CPU E5-2667 v3 @ 3.20 GHz (8 cores, 16 threads per socket); 4×16 Go @ 2400 MHz RAM. Code was run on Python 3.6 on Docker (18.09.0) containers based on the official python 3.6-stretch container, available on ETFL GitHub.

## Supporting information

**S1 Nondimensionalization. Derivation and formulation.** Details on the variables and constraint transformations to scale the model.

**S2 Example EP constraint matrix. Representation of the constraint matrix of the EP for a vETFL of iJO1366.** Colored cells represent non-zero blocks. Uncolored cells are zero blocks.

**S3 SOP for creating an ETFL model. Tips and Prerequisites.** List of required and optional inputs to transform a COBRA model into an ETFL model

## Acknowledgements

The authors would like to thank Kaycie Butler for her valuable input on the wording and structure of this manuscript. This work has received funding from the European Union’s Horizon 2020 Research and Innovation Programme under the Marie Sklodowska-Curie grant agreement No 722287.

