## Supplementary Information S1 for "ETFL: A formulation for flux balance models accounting for expression, thermodynamics, and resource allocation constraints" (from another version of this preprint: version 1, 2019-03-28)

### Nondimensional Scaling

A critical issue in the formulation of this problem is the different orders of magnitude the variables belong to. Fluxes are typically between $10^{-3} - 10^{1}\ mmol.gDW^{-1}.h^{-1}$. Protein concentrations are around $10^{-6} - 10^{-3}\ mmol.gDW^{-1}$, and mRNA concentrations $10^{-10} - 10^{-6}\ mmol.gDW^{-1}$. The relationship between these scales is given by the catalytic rates of enzymes and expression machinery, which span $10^{3} - 10^{6}\ h^{-1}$. As a consequence, the constraint matrix becomes ill-conditioned and the solver has to operate close to, or sometimes beyond, their maximal solving accuracy (usually around $10^{-9}$ for commercial solvers such as ILOG CPLEX or Gurobi)

In order to circumvent these limitations, we operate a scaling of the EP which will reduce the numerical difficulty of the problem.

In particular, we consider nondimensionalization by upper bound as a method, which will also allow to reduce the effective range of the variables seen by the solver.

Page 2 details the full bilinear EP formulation, before and after scaling. Page 3 introduces nondimensionalization constants. Page 4 shows the relationship between original variables and scaled variables. Page 5 summarizes upper bounds used for nondimensionalization, and the nondimensionalization variables.

| ***Initial*** | ***Scaled*** | |
|---|---|---|
| $S \cdot v = 0$ | | *(FBA)* |
| *Catalytic constraints* | | |
| $v_j^f - k_{cat,f}^j E_j \leq 0$ | $\tilde{v}_j^f - \tilde{E}_j \leq 0$ | *(FC)*$_j$ |
| $v_j^b - k_{cat,b}^j E_j \leq 0$ | $\tilde{v}_j^b - \tilde{E}_j \leq 0$ | *(BC)*$_j$ |
| *Macromolecule mass balances* | | |
| $v_j^{asm} - v_j^{deg} - \mu * E_j = 0$ | $\tilde{v}_j^{asm} - \tilde{v}_j^{deg} - \frac{\mu}{k_{deg}^j} * \tilde{E}_j = 0$ | *(EB)*$_j$ |
| $v_l^{tcr} - v_l^{deg} - \mu * F_l = 0$ | $\frac{1}{k_{deg}^l F_l^{ub}} \cdot \frac{k_{cat}^{RNAP} P_l^{ub}}{L_l^{nt}} \cdot \tilde{v}_l^{tcr} - \tilde{v}_l^{deg} - \frac{\mu}{k_{deg}^l} * \tilde{F}_l = 0$ | *(MB)*$_l$ |
| $v_l^{tsl} - \sum_j \eta_l^j \cdot v_j^{asm} = 0$ | $\tilde{v}_l^{tsl} - \sum_j \eta_l^j \cdot \frac{L_l^{aa}}{k_{cat}^{rib} R_l^{ub}} \cdot k_{deg}^j E_j^{ub} \cdot \tilde{v}_j^{asm} = 0$ | *(PB)*$_l$ |
| $-v_{aa_i}^{charging} + \sum_l \eta_{aa_i}^l \cdot v_l^{tsl} - \mu * T_{aa_i}^u = 0$ | $-\frac{1}{\bar{\mu}\, T_{aa_i}^{ub}} v_{aa_i}^{charging} + \sum_l \eta_{aa_i}^l \cdot \frac{1}{\bar{\mu}\, T_{aa_i}^{ub}} \cdot \frac{k^{rib} R_l^{ub}}{L_l^{aa}} \cdot \tilde{v}_l^{tsl} - \frac{\mu}{\bar{\mu}} * \tilde{T}_{aa_i}^u = 0$ | *(TB)*$_{aa,i,u}$ |
| $v_{aa_i}^{charging} - \sum_l \eta_{aa_i}^l \cdot v_l^{tsl} - \mu * T_{aa_i}^c = 0$ | $\frac{1}{\bar{\mu}\, T_{aa_i}^{ub}} v_{aa_i}^{charging} - \sum_l \eta_{aa_i}^l \cdot \frac{1}{\bar{\mu}\, T_{aa_i}^{ub}} \cdot \frac{k^{rib} R_l^{ub}}{L_l^{aa}} \cdot \tilde{v}_l^{tsl} - \frac{\mu}{\bar{\mu}} * \tilde{T}_{aa_i}^u = 0$ | *(TB)*$_{aa,i,c}$ |
| $v_{rRNA_l}^{tcr} - v_{rib}^{asm} = 0$ | $\frac{1}{k_{deg}^j E_j^{ub}} \cdot \frac{k_{cat}^{RNAP} P_l^{ub}}{L_l^{nt}} \cdot \tilde{v}_{rRNA_l}^{tcr} - \tilde{v}_{rib}^{asm} = 0$ | *(RB)*$_{rRNA,l}$ |
| *Macromolecule degradation* | | |
| $v_j^{deg} - k_{deg}^j \cdot E_j = 0$ | $\tilde{v}_j^{deg} - \tilde{E}_j = 0$ | *(ED)*$_j$ |
| $v_l^{deg} - k_{deg}^l \cdot F_l = 0$ | $\tilde{v}_l^{deg} - \tilde{F}_l = 0$ | *(MD)*$_l$ |
| *Transcript./translation catalytic constraints* | | |
| $v_l^{tsl} - \frac{k_{cat}^{rib}}{L_l^{aa}} R_l \leq 0$ | $\tilde{v}_l^{tsl} - \tilde{R}_l \leq 0$ | *(TR2)*$_l$ |
| $v_l^{tcr} - \frac{k_{cat}^{RNAP}}{L_l^{nt}} P_l \leq 0$ | $\tilde{v}_l^{tcr} - \tilde{P}_l \leq 0$ | *(TR1)*$_l$ |
| *Expression coupling* | | |
| $R_l - \frac{L_l^{nt}}{L_{rib}^{nt}} F_l \leq 0$ | $\tilde{R}_l - \frac{L_l^{nt}}{L_{rib}^{nt}} \frac{F_l^{ub}}{R_l^{ub}} \tilde{F}_l \leq 0$ | *(EX)*$_l$ |
| *Capacity constraints* | | |
| $R_F - (1-\rho) E_{rib} = 0$ | $\tilde{R}_F - (1-\rho) \tilde{E}_{rib} = 0$ | *(RR)* |
| $\sum_l R_l + R_F - E_{rib} = 0$ | $\sum_l \frac{R_l^{ub}}{E_{rib}^{ub}} \tilde{R}_l + \tilde{R}_F - \tilde{E}_{rib} = 0$ | *(TC2)* |
| $\sum_l P_l - E_{RNAP} = 0$ | $\sum_l \frac{P_l^{ub}}{E_{RNAP}^{ub}} P_l - \tilde{E}_{RNAP} = 0$ | *(TC1)* |

| ***Initial*** | ***Scaled*** | |
|---|---|---|
| $S \cdot v = 0$ | | *(FBA)* |
| *Catalytic constraints* | | |
| $v_j^f - k_{cat,f}^j E_j \le 0$ | $\tilde{v}_j^f - \tilde{E}_j \le 0$ | *(FC)*$_j$ |
| $v_j^b - k_{cat,b}^j E_j \le 0$ | $\tilde{v}_j^b - \tilde{E}_j \le 0$ | *(BC)*$_j$ |
| *Macromolecule mass balances* | | |
| $v_j^{asm} - v_j^{deg} - \mu * E_j = 0$ | $\tilde{v}_j^{asm} - \tilde{v}_j^{deg} - \alpha_j \cdot \mu * \tilde{E}_j = 0$ | *(EB)*$_j$ |
| $v_l^{tcr} - v_l^{deg} - \mu * F_l = 0$ | $\gamma_l \cdot \tilde{v}_l^{tcr} - \tilde{v}_l^{deg} - \beta_j \cdot \mu * \tilde{F}_l = 0$ | *(MB)*$_l$ |
| $v_l^{tsl} - \sum_j \eta_l^j \cdot v_j^{asm} = 0$ | $\tilde{v}_l^{tsl} - \sum_j \eta_l^j \cdot \delta_l^j \cdot \tilde{v}_j^{asm} = 0$ | *(PB)*$_l$ |
| $-v_{aa_i}^{charging} + \sum_l \eta_{aa_i}^l \cdot v_l^{tsl} - \mu * T_{aa_i}^u = 0$ | $-\frac{1}{\bar{\mu}\, T_{aa_i}^{ub}} v_{aa_i}^{charging} + \sum_l \eta_{aa_i}^l \cdot \tau_l \cdot \tilde{v}_l^{tsl} - \frac{\mu}{\bar{\mu}} * \tilde{T}_{aa_i}^u = 0$ | *(TB)*$_{aa,i,u}$ |
| $v_{aa_i}^{charging} - \sum_l \eta_{aa_i}^l \cdot v_l^{tsl} - \mu * T_{aa_i}^c = 0$ | $\frac{1}{\bar{\mu}\, T_{aa_i}^{ub}} v_{aa_i}^{charging} - \sum_l \eta_{aa_i}^l \cdot \tau_l \cdot \tilde{v}_l^{tsl} - \frac{\mu}{\bar{\mu}} * \tilde{T}_{aa_i}^u = 0$ | *(TB)*$_{aa,i,c}$ |
| $v_{rRNA_l}^{tcr} - v_{rib}^{asm} = 0$ | $\gamma_l \cdot \tilde{v}_{rRNA_l}^{tcr} - \tilde{v}_{rib}^{asm} = 0$ | *(RB)*$_{rRNA,l}$ |
| *Macromolecule degradation* | | |
| $v_j^{deg} - k_{deg}^j \cdot E_j = 0$ | $\tilde{v}_j^{deg} - \tilde{E}_j = 0$ | *(ED)*$_j$ |
| $v_l^{deg} - k_{deg}^l \cdot F_l = 0$ | $\tilde{v}_l^{deg} - \tilde{F}_l = 0$ | *(MD)*$_l$ |
| *Transcript./translation catalytic constraints* | | |
| $v_l^{tsl} - \frac{k_{cat}^{rib}}{L_l^{aa}} R_l \le 0$ | $\tilde{v}_l^{tsl} - \tilde{R}_l \le 0$ | *(TR2)*$_l$ |
| $v_l^{tcr} - \frac{k_{cat}^{RNAP}}{L_l^{nt}} P_l \le 0$ | $\tilde{v}_l^{tcr} - \tilde{P}_l \le 0$ | *(TR1)*$_l$ |
| *Expression coupling* | | |
| $R_l - \frac{L_l^{nt}}{L_{rib}^{nt}} F_l \le 0$ | $\tilde{R}_l - \lambda_l \cdot \tilde{F}_l \le 0$ | *(EX)*$_l$ |
| *Capacity constraints* | | |
| $R_F - (1 - \rho)E_{rib} = 0$ | $\tilde{R}_F - (1 - \rho)\tilde{E}_{rib} = 0$ | *(RR)* |
| $\sum_l R_l + R_F - E_{rib} = 0$ | $\sum_l \rho_l \cdot \tilde{R}_l + \tilde{R}_F - \tilde{E}_{rib} = 0$ | *(TC2)* |
| $\sum_l P_l - E_{RNAP} = 0$ | $\sum_l \pi_l \cdot P_l - \tilde{E}_{RNAP} = 0$ | *(TC1)* |

| ***Variable*** | ***Nondimensionalization factor*** | ***Scaled variable*** |
|---|---|---|
| $X$ | $\alpha$ | $\tilde{X} = \dfrac{X}{\alpha}$ |
| $E_j$ | $E_j^{ub}$ | $\tilde{E}_j = \dfrac{E_j}{E_j^{ub}}$ |
| $F_l$ | $F_l^{ub}$ | $\tilde{F}_l = \dfrac{F_l}{F_l^{ub}}$ |
| $R_l$ | $R_l^{ub}$ | $\tilde{R}_l = \dfrac{R_l}{R_l^{ub}}$ |
| $R_F$ | $E_{rib}^{ub}$ | $\tilde{R}_F = \dfrac{R_F}{E_{rib}^{ub}}$ |
| $P_l$ | $P_l^{ub}$ | $\tilde{P}_l = \dfrac{P_l}{P_l^{ub}}$ |
| $T_{aa_i}^u$ | $T_{aa_i}^{ub}$ | $\tilde{T}_{aa_i}^u = \dfrac{T_{aa_i}^u}{T_{aa_i}^{ub}}$ |
| $T_{aa_i}^c$ | $T_{aa_i}^{ub}$ | $\tilde{T}_{aa_i}^c = \dfrac{T_{aa_i}^c}{T_{aa_i}^{ub}}$ |
| $v_j^f$ | $k_{cat,f}^j E_j^{ub}$ | $\tilde{v}_j^f = \dfrac{v_j^f}{k_{cat,f}^j E_j^{ub}}$ |
| $v_j^b$ | $k_{cat,b}^j E_j^{ub}$ | $\tilde{v}_j^b = \dfrac{v_j^b}{k_{cat,b}^j E_j^{ub}}$ |
| $v_j^{deg}$ | $k_{deg}^j E_j^{ub}$ | $\tilde{v}_j^{deg} = \dfrac{v_j^{deg}}{k_{deg}^j E_j^{ub}}$ |
| $v_l^{deg}$ | $k_{deg}^l F_l^{ub}$ | $\tilde{v}_l^{deg} = \dfrac{v_l^{deg}}{k_{deg}^l F_l^{ub}}$ |
| $v_l^{tsl}$ | $\dfrac{k^{rib}}{L_l^{aa}} R_l^{ub}$ | $\tilde{v}_l^{tsl} = \dfrac{L_l^{aa}}{k_{cat}^{rib} R_l^{ub}} \cdot v_l^{tsl}$ |
| $v_l^{tcr}$ | $\dfrac{k_{cat}^{RNAP}}{L_l^{nt}} P_l^{ub}$ | $\tilde{v}_l^{tcr} = \dfrac{L_l^{nt}}{k_{cat}^{RNAP} P_l^{ub}} \cdot v_l^{tcr}$ |
| $v_j^{asm}$ | $k_{deg}^j E_j^{ub}$ | $\tilde{v}_j^{asm} = \dfrac{v_j^{asm}}{k_{deg}^j E_j^{ub}}$ |

| *Upper bound* | *(Approximate) Value* | *Order of magnitude* | *Unit* |
|---|---|---|---|
| $E_j^{ub}$ | $\frac{1}{MW\left(Enz_j\right)} \cdot 10^3$ | $10^{-2} \cdot 10^3 = 10^1$ | $\frac{[g.gDW^{-1}]}{[g.mol^{-1}]}\left[\frac{mmol}{mol}\right] = mmol.gDW^{-1}$ |
| $F_l^{ub}$ | $\frac{1}{MW(mRNA_l)} \cdot 10^3$ | $10^{-2} \cdot 10^3 = 10^1$ | $mmol.gDW^{-1}$ |
| $R_l^{ub}$ | $\frac{1}{MW(Rib)} \cdot 10^3$ | $10^{-6} \cdot 10^3 = 10^{-3}$ | $mmol.gDW^{-1}$ |
| $P_l^{ub}$ | $\frac{1}{MW(RNAP)} \cdot 10^3$ | $10^{-5} \cdot 10^3 = 10^{-2}$ | $mmol.gDW^{-1}$ |
| $T_{aa_i}^{ub}$ | $\frac{1}{MW\left(tRNA_{aa_i}\right)} \cdot 10^3$ | $10^{-2} \cdot 10^3 = 10^1$ | $mmol.gDW^{-1}$ |

| *Nondimensionalization term* | *Expression* | *Order of magnitude* |
|---|---|---|
| $\alpha_j$ | $\frac{\mu}{k_{deg}^j}$ | |
| $\beta_l$ | $\frac{\mu}{k_{deg}^l}$ | |
| $\gamma_l$ | $\frac{1}{k_{deg}^l F_l^{ub}} \cdot \frac{k_{cat}^{RNAP} P_l^{ub}}{L_l^{nt}}$ | |
| $\delta_l^j$ | $\frac{L_l^{aa}}{k_{cat}^{rib} R_l^{ub}} \cdot k_{deg}^j E_j^{ub}$ | |
| $\tau_l^{aa_i}$ | $\frac{1}{\bar{\mu}\, T_{aa_i}^{ub}} \cdot \frac{k^{rib} R_l^{ub}}{L_l^{aa}}$ | |
| $\lambda_l$ | $\frac{L_l^{nt}}{L_{rib}^{nt}} \frac{F_l^{ub}}{R_l^{ub}}$ | |
| $\rho_l$ | $\frac{R_l^{ub}}{E_{rib}^{ub}}$ | |
| $\pi_l$ | $\frac{P_l^{ub}}{E_{RNAP}^{ub}}$ | |
