## Supplementary Information S3 for "ETFL: A formulation for flux balance models accounting for expression, thermodynamics, and resource allocation constraints" (from another version of this preprint: version 1, 2019-03-28)

### ETFL: Supplementary Information: SOP for creating an ETFL model

#### Summary checklist

Here is a summarized checklist of the material needed to turn a COBRA model into ETFL:

- A working installation of ETFL
- A Cobra model with:
  - Gene identifiers (IDs)
  - All nucleotides triphosphates(NTPs), deoxynucleotides triphosphate(dNTP), nucleotides diphosphate (NMP), aminoacids.
  - (Optional) Gene reaction rules
- Gene sequences indexed by their gene IDs
- Peptide stoichiometry of enzymes
- Enzyme assigments per reaction.
- Enzyme catalytic rate constants:
  - Forward
  - (Optional) Reverse
- Enzyme degradation rate constants
- mRNA degradation rate constants
- (Optional) Free ribosomes ratio
- (Optional) Free RNA Polymerase ratio
- (Optional) GC-content and length of the genome
- (Optional) Average aminoacid abundances
- (Optional) Average NTP abundances
- (Optional) Average mRNA length
- (Optional) Avergae peptide length
- (Optional) Growth-dependant mRNA, peptide, and DNA mass ratios.

### 1 Setup

#### 1.1 Prerequisites

Make sure you have `Git` installed. Since ETFL is built upon pyTFA [1], we will clone both repositories. In a folder of your choice, download the source code from our repositories:

```
git clone https://github.com/EPFL-LCSB/pytfa
git clone https://github.com/EPFL-LCSB/etfl
# -- OR --
git clone https://gitlab.com/EPFL-LCSB/pytfa
git clone https://gitlab.com/EPFL-LCSB/etfl
```

#### 1.2 Docker container (recommended)

We recommend the use of Docker containers as they provide a standardized, controlled and reproducible environment. The ETFL Docker is built upon the pyTFA Docker image. We recommend building it yourself as it is where your solvers can be installed.

##### 1.2.1 Downloading Docker

If Docker is not yet installed on your machine, you can get it from [here]

##### 1.2.2 Building and running the Docker container

```
# Build the pyTFA docker
cd pytfa/docker && . build
# Build and run the ETFL docker
cd ../../etfl/docker
. build
. run
```

##### 1.2.3 Solvers

For installing the solvers, please refer to the pyTFA documentation

#### 1.3 Python environment

Alternatively, you can install ETFL using either:

```
pip install -r etfl/requiremetns.txt
# -- OR --
pip install etfl
```

Make sure your solvers are also installed in the same environment if you are using a `virtualenv` or `pyenv`.

### 2 From COBRA to ETFL

ETFL models can be generated fairly easily from a COBRA model. In the following subsections, we detail the required information to add expression constraints to a COBRA model and turn it into an ETFL model.

#### 2.1 Constraint-based model

You will need to start with a COBRA model including the following information:

- Genes and their gene ID (necessary to retrieve gene sequences)
- (Optional) Gene-protein rules: These are used to make approximated enzymes if peptide information is not enough

Additionally, you will need to build a dictionnary of essential metabolites required in the model. It should follow this example structure (all fields mandatory):

```
dict(atp='atp_c',
     adp='adp_c',
     amp='amp_c',
     gtp='gtp_c',
     gdp='gdp_c',
     pi ='pi_c' ,
     ppi='ppi_c',
     h2o='h2o_c',
     h  ='h_c'  )
```

A dictionnary of RNA NTPs, DNA dNTPS, and aminoacids is also required, of the type:

```
aa_dict = {'A': 'ala__L_c',
           # ...
           'V': 'val__L_c', }

rna_nucleotides = {
                'u': 'utp_c',
                # ...
                'c': 'ctp_c'}

rna_nucleotides_mp = {
                'u': 'ump_c',
                # ...
                'c': 'cmp_c'}

dna_nucleotides = {
                't': 'dttp_c',
                # ...
                'c': 'dctp_c'}
```

#### 2.2 From genes to peptides

In order to build the transcription and translation, it is necessary to provide ETFL with gene deoxynucleotide sequences. These will be automatically transcribed in RNA sequences and then translated into aminoacid peptide sequences. They must be fed to the function `model.add_nucleotides_sequences` in a dict-like object, indexed by gene IDs (`model.genes.mygene.id` property in COBRA).

We suggest the following sources for obtaining such information:

- KEGG Genes
- NCBI Gene DB

- MetaCyc Gene Search

ETFL will automatically synthesize the correct peptides from the nucleotides sequences. This is based on the Biopython package's `transcribe` and `translate` functions [2].

For each enzyme created by transcription, a degradation rate constant must be specified. These can be obtained through literature search, or using an average value.

##### 2.3 From peptides to enzymes

A key part of the expression modeling is to properly represent the assembly of enzymes from peptides. For each enzyme of the model, a stoichiometry of the peptides necessary for its assembly is needed. These are stored as dictionnaries in the `Enzyme.composition` property under a form similar to :

```
>>> enzyme.composition
{'b2868': 1, 'b2866': 1, 'b2867': 1}
```

The keys match the IDs of genes coding for the peptide, and the value represent the stoichiometry of the peptide in the enzyme. These can be obtained from litterature search or specialized databases. In particular, we used for this paper the Metacyc/Biocyc database [3, 4], using specialised SmartTables queries [5].

```
html-sort-ascending(
    html-table-headers (
        [(f,genes,(protein-to-components f)):
            f<-ECOLI^^Protein-Complexes,genes := (enzyme-to-genes f)
        ],
        ("Product Name", "Genes", "Component coefficients")),
    1)
```

##### 2.4 From enzymes back to the metabolism

Lastly, the enzymes must be assigned reactions and catalytic rate constants. Several enzymes can catalyze the same reactions. COBRA models can take this into account differently, usually having either (i) multiple reactions with a simple gene reaction rule; or (ii) one unique reaction with several isozymes in the gene reaction rule. Although not often applied consistently within the same model, these two formalisms are equivalent, and their ETFL counterparts will also behave equivalently.

For each enzyme, the information needed is the (forward) catalytic rate constant $k_{cat}^{+}$, facultatively the reverse catalytic rate constant $k_{cat}^{-}$ (set equal to $k_{cat}^{+}$ if none is given), and a degradation rate constant.

This is done by calling the function `model.add_enzymatic_coupling(coupling_dict)` where `coupling_dict` is a dict-like object with reaction IDs as keys and a list of enxyme objects as values:

```
coupling_dict = {
    #...
    'AB6PGH': [ <Enzyme AB6PGH_G495_MONOMER at 0x7ff00e0f1b38>],
    'ABTA'  : [ <Enzyme ABTA_GABATRANSAM at 0x7ff00e0fda90>,
                <Enzyme ABTA_G6646 at 0x7ff00e0fd4e0>],
    'ACALD' : [ <Enzyme ACALD_MHPF at 0x7ff00e0fdcf8>],
    #...
    }
```

The catalytic rate constants can be obtained from several databases, such as:

- Rhea
- BRENDA
- SabioRK
- Uniprot

Several enzymes can be assigned to a reaction. ETFL will try to match the gene reaction rule isozymes to the supplied enzymes. If the gene reaction rule shows several isozymes while only one enzyme is supplied, the enzyme can be replicated to match the number of isozymes in the gene reaction rule.

Given a reaction in the model, if no enzyme is supplied but the reaction possesses a gene reaction rule, it is possible to infer an enzyme from it. The rule expression is expanded, and each term seprated a by an `OR` boolean operator is interpreted as an isozyme, while terms separated by an `AND` boolean operators are interpreted as unit peptide stoichiometric requirements. The enzyme is then assigned an average catalytic rate constant and degradation rate constant.

### 3 Growth-dependant parameters

Accounting for growth-dependent RNA and protein content requires additional information. In particular:

- GC-content and length of the genome
- Average aminoacid abundances
- Average NTP abundances
- Average mRNA length
- Avergae peptide length
- Growth-dependant mRNA, peptide, and DNA mass ratios.

These values are usually obtained through litterature search. All of the last three ratios are optional, although using none defeats the purpose of accounting for growth-dependant parameters.

### 4 Additional documentation

#### 4.1 Example

We encourage the reader to look at the script used to generate the models with which the paper's results were generated, available in `etfl/tutorials/helper_gen_models.py`. The data it takes in input has been generated in `etfl/etfl/data/ecoli.py`. These are good examples to start from in order to make a custom ETFL from a different COBRA model.

### Acknowledgments

This work has received funding from the European Union's Horizon 2020 research and innovation programme under the Marie Skłodowska-Curie grant agreement No 722287.
